# XAMPGen: An Interpretable Super Intelligence Pipeline For Novel Antimicrobial Peptide Discovery

**DOI:** 10.64898/2026.11.03.764041

**Authors:** Kinjal Mondal, Sarala Padi, Navleen Kaur, Wright Makambi, David P. Hoogerheide, Frank Heinrich, Mihaela Mihailescu, Jeffery B. Klauda, Antonio Cardone

## Abstract

Antimicrobial peptides (AMPs) are short peptides with a broad spectrum of activity against bacterial pathogens, representing a promising avenue for combating the growing threat of antibiotic resistance. This study focuses on two main components: 1) generative super intelligence (GenSI) and 2) utilizing explainable super intelligence (XSI). We fine-tuned ProGen, a decoder-only protein language model, on known AMPs to generate novel peptide sequences with strong antimicrobial properties. Generated candidates were screened with an AMP classifier and filtered based on predicted minimum inhibitory concentration (MIC), along with human insights into AMP structure and activity in bacterial membranes. A final set of peptides was selected for synthesis and experimental testing. Biological assays confirmed antimicrobial activity in all generated and tested peptides, while fluorescence and circular dichroism measurements characterized membrane binding and peptide structure. We then applied XSI techniques to interpret the generative model and derive biological insights. Specifically, layer probing was used to identify cornerstone layers that encode functionally critical information. Self-attention analysis within these layers revealed residue-level interdependencies and patterns underlying antimicrobial functions, which we evaluated against well-characterized AMPs from the literature. We further examined associations between amino acid sequence patterns and binding properties in bacterial membrane mimics, relating these observations to biological activities in cell-based assays. These analyses provide insight into the model’s decision-making process. This builds confidence that the learned amino acid interdependencies have physico-chemical significance, as applied to AMPs, and may facilitate the discovery of novel AMPs with therapeutic potential. This work demonstrates that interpretable generative language models can support rational AMP discovery.

## Introduction

Antimicrobial resistance (AMR) represents a significant global challenge, with millions of deaths attributed to this growing crisis [1]. A primary driver of AMR is that conventional antibiotics are typically small molecules—comprising only 15 to 60 heavy atoms (30 to 120 total atoms) and have narrow, specific mechanisms of action, making it relatively easy for bacteria to develop resistance through targeted genetic mutations [2]. Antimicrobial peptides [3, 4] (AMPs) are short cationic sequences 10-50 amino acids long that provide an alternative solution to combat bacterial infection. AMPs exhibit various mechanisms of action; however, many primarily target bacterial cell membranes, making it more challenging for bacteria to develop resistance. As such, AMPs have emerged as a viable alternative to traditional antibiotics [5].

Despite their therapeutic potential, the number of known AMPs remains limited. The vast peptide sequence space further complicates AMP design, making experimental trial-and-error approaches impractical. Many AMPs adopt secondary structures upon interacting with lipid membranes [6]. These structural transitions are thought to be important in their antimicrobial activity [6]. However, their structures in this environment remain poorly characterized, which makes rational design difficult [7, 8]. Computational methods, specifically recent advances in machine learning, have advanced research that can be used to generate AMP sequences, and predict their properties and structures [9, 10, 11].

The advent of protein language models (PLMs) [12, 13, 14, 15] has marked a new era in computational biology. Trained in millions of sequences, these models encode high quality representations of the diverse protein space and learn evolutionary, structural, and functional information through their training. In particular, ProGen [13], a decoder-only language model, has been shown to generate functional protein families after fine-tuning. Similarly, encoder-only language models such as Evolutionary Scale Modeling (ESM) [14, 15] excel in protein sequence understanding and representation, which can then be used for downstream tasks such as protein function/structure prediction.

Recently, there has been a huge interest in applying these pretrained PLMs towards designing AMPs as well as predicting their properties. Among these, the last couple of years has seen several such models based on the Generative Pre-trained Transformer (GPT) architecture, such as ProteoGPT [16], AMPDesigner [17], BroadAMPGPT [18], AMPCLGPT [19], and reinforcement-learning driven GPT [20], prompt-tuning based on ProtGPT2 [21] used for designing and generating AMPs. A particular challenge in adapting these large language models is the significant computational cost of full fine-tuning. These models contain millions to billions of parameters that require a lot of memory and compute resources. To tackle these challenges, several parameter-efficient fine-tuning (PEFT) methods have been developed [22, 23] to fine-tune LLMs. In this work, we adopt the LoRA adaptation in the ProGen model for the generation of antimicrobial peptides.

Despite the remarkable progress in the use of PLMs for the design of AMPs for the last couple of years, the process of AMP design has remained largely a black box. These models offer limited insight into the sequence characteristics and inter-residue relationships learned through their self-attention layers or how these features influence their predictions. This opacity is a significant barrier in the overall context of peptide design, where understanding the internal basis of a model’s predictions is essential for building confidence in the model and for guiding rational downstream optimization. To address these challenges, we develop XAMPGen (Explainable AMP Generation), an SI method built upon techniques from Explainable Super Intelligence (XSI) [24] to explain the inner workings for sequence generation of our fine-tuned ProGen model.

We generate candidate AMPs using our fine-tuned ProGen model, which are then filtered using (1) a classifier and (2) a regressor built on the pretrained ESM [15] model. Finally, we select sequences on the basis of length, hydrophobicity, and synthesis costs. Fifteen peptides were selected for experimental measurements, starting with bacterial susceptibility assays for determining their Minimal Inhibitory Concentration (MIC). We further use fluorescence and circular dichroism experiments to study the interactions between these peptides and lipid membranes mimicking bacterial membranes — the primary target of the majority of known AMPs.

Next, we move to the XSI core of our method. Within ProGen, each transformer layer is a successive computational stage that updates an internal numerical representation of the sequence generated so far. We examine these representations across layers to identify where AMP-related properties can be predicted most accurately. On the basis of layer probing, we then select a *cornerstone layer* — the layer whose internal representation carries the most task-relevant information, and which we use for all the biological claims that follow. We introduce a normalization technique for the self-attention maps derived by the model to interpret the hidden correlations between the various tokens, thereby deriving useful biological insight. We use our interpretable method to analyze known AMPs as well as AMPs previously measured in our lab [7] to explain their mechanisms. Finally, we use our XSI method and the experimental observables to provide rational explanations linking the XSI observations and the experimental measurements for the newly generated peptides. A schematic of our pipeline is given in **Fig 1**.

**Figure 1.**
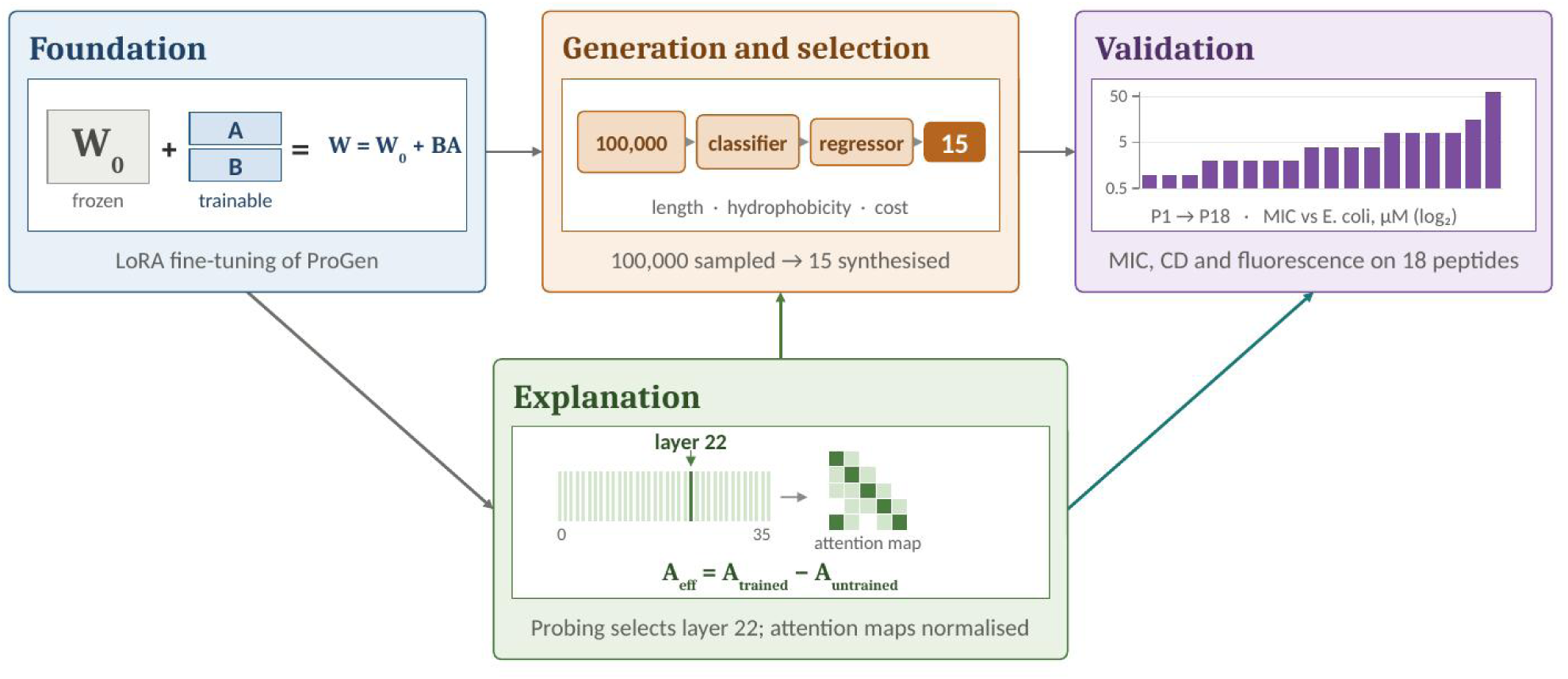
An overview of our generation–validation–explanation components of the XAMPGen pipeline for AMP design.

## Methods and Materials

### Fine-tuning of ProGen

ProGen [13] is a GPT-based decoder-only PLM trained on 280 million protein sequences drawn from UniParc [25], UniProtKB [26], Pfam [27], and NCBI [28], and conditioned on the functional metadata aggregated from those same databases. This model follows the causal (masked) self-attention architecture, where each token only attends to preceding tokens. Formally, the model is trained to minimize the following negative log-likelihood over the dataset

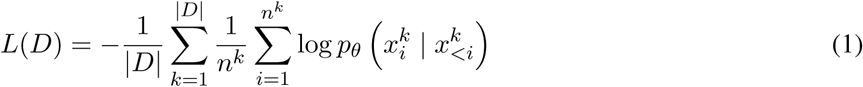

where *k* indexes a sequence in the training set, *i* indexes a token position within that sequence, and *L*(*D*) represents the total loss over the dataset, |*D*| denotes the total number of sequences in the dataset or the training batch, and *n_k_* represents the number of tokens in the *k*-th sequence. The term 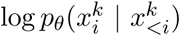 represents the logarithmic conditional probability of the *i*-th token given all preceding tokens in the *k*-th sequence under model parameters *θ*. The token sequence is written as *x* = [*c, a*], where 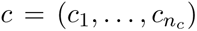 contains functional control tags and 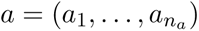 contains amino acid tokens. A control tag specifies a desired property, such as “antimicrobial”; an amino acid token identifies a residue, such as alanine (A) or lysine (K). Each tag or residue is assigned a vocabulary index that the model maps to a learned embedding vector. Thus, “antimicrobial” is a functional control tag represented by a token in the model vocabulary. ProGen is trained in two stages, which use these control tags differently. In pretraining, the model is trained across many protein families at once, with the control tags identifying the family and functional properties of each sequence; it therefore learns broad sequence patterns and how those patterns depend on the tags. In fine-tuning, we train the model on a single family under a single tag — antimicrobial peptides in our case — so that the general sequence knowledge acquired during pretraining is specialized to that family. The decoder-only architecture of the transformer is trained through the following causal self-attention mechanism. First, for a sequence of L tokens, we project them into a learnable embedding space *X* ∈ ℝ*^L×d^*, which is transformed into three different projections, query (Q), Key (K), and Value (V) through the learnable matrices *W_Q_*, *W_K_*, and *W_V_* :

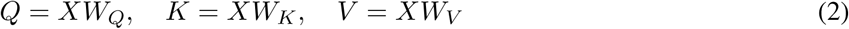

This decoupled representation allows each token to dynamically express what information it seeks (*Q*), what context it offers (*K*), and the actual core content it communicates (*V*). Finally, a learnable self-attention matrix is computed, which provides information regarding the relative importance of one token to another. The self-attention matrix *A* ∈ ℝ*^L×L^* is computed via scaled dot-product attention:

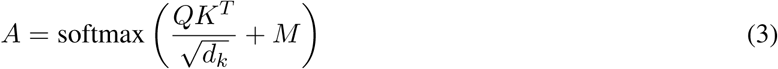

where *d_k_* represents the dimension of the key vector. The matrix *M* enforces the causal property, defined as *M_ij_*= 0 if *j ≤ i* and *M_ij_* = −∞ otherwise. Thus, at position *i* the model can use the current and preceding tokens, but cannot gather context from future positions *j > i*.

Full fine-tuning updates all pretrained model parameters on the target dataset and is expensive since they contain millions or billions of parameters, and optimizing them requires large computational resources and time. Thereby, several low-cost and memory-efficient methods have been developed [22, 23] to fine-tune LLM’s. In our paper, we adapt an efficient fine-tuning technique called Low-Rank Adaptation (LoRA). In LoRA, we freeze the original model parameters (*W*_0_) and instead train two smaller, lower-rank matrices, A and B. The overall adapted weight matrix is then written as

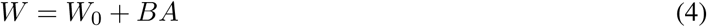

We take the “antimicrobial” control tag present in the functional keywords of the training data for ProGen and append it to the antimicrobial peptide datatset collected by us. Post-training, this tag serves as a specific prompt to conditionally generate novel antimicrobial peptides.

### The Generation and Filtering pipelines

Having fine-tuned ProGen on known AMPs, we next generated new peptide sequences and filtered them down to a final set for synthesis. To generate peptides, we used a top-p sampling criterion (p=0.9). At each generation step, the sampling vocabulary was limited to the subset of tokens whose cumulative sampling probability exceeded p, thereby truncating the low-probability tail in order to design more functional peptides.

To prepare peptides for experimental design, we developed SI tools that predict whether a peptide is antimicrobial and estimate the MIC of AMP sequences. To do this, we trained a separate classifier and regressor, both based on the ESM model. The ESM [15, 14] model differs from the decoder-only ProGen in that it is an encoder-only model trained through a Masked Language Modeling (MLM) objective on protein sequences, thereby giving us an overall representation on these protein sequences, making it exceptionally suited for predictive tasks instead of generative ones. For training the classifier and the regressor, we take the model, extract the representations from the last layer, do a mean pooling and fit a linear layer on top of them to train the AMP classifier and the MIC regressor.

Once trained, the generated sequences were routed through our trained classifier and regressor to finalize a list of peptides. We further narrowed the selection by prioritizing sequences with optimal length (10–30 residues) and favorable hydrophobicity, ensuring the peptides were sufficiently water-soluble for reliable biophysical characterization. Additionally, we targeted candidate sequences located close within the ESM representation space to a highly potent, natural AMP (Tilapia Piscidin 4), for ease of comparing with biophysical measurements. Once we have generated several peptides and measured their MICs, we use our XSI method, described in the next subsection, to gain further insights into their sequence characteristics and measurable and interpretable physico-chemical properties. Here, we focus on measurements of biological activity with in vitro, antibacterial assays (MIC determination) and peptide binding affinities and structures at lipid membranes - the primary target of most AMPs in bacteria.

### Interpretation for Fine-tuned ProGen

Having described how candidate peptides are generated and filtered, we now turn to the interpretation of the model itself. To understand how the model generates antimicrobial peptides and to interpret its hidden layers, we adopted the following XSI strategies. We probe the representations at every layer on two sequence-level tasks, AMP classification and MIC regression (as explained below), and we separately extract and normalize the causal attention maps of every layer, from which we compute an attention-derived proxy for helicity. Taken together, these three measurements single out one *cornerstone layer*, and the biological analysis that follows is carried out on that layer.

Layer probing aims to identify the layers that encode the most robust task-specific information. In this approach, we take the representations at each layer and use them for the following tasks: (1) a binary classification between AMPs and non-AMPs and (2) a regression task for MIC activity prediction. For the third task, we analyze the self-attention heatmaps and compute approximate helicities from them for each layer, which we then compare with the helicities measured experimentally by CD spectroscopy. The detailed approach is described below.

The self-attention maps help us understand how the model captures cross-token interactions, representing underlying biochemical correlations between amino acid residues. However, due to the causal architectural bias in ProGen (one token only attends to preceding tokens), the tokens in an early row of the self attention matrix do not see the same number of tokens as those in later rows. Thereby, attention values in earlier rows might be much higher than the values in the later rows due to the fact that the earlier rows are distributing probability over fewer tokens. To isolate actual inter-residue biological correlations from this architectural bias, we introduce a normalization technique. We define an untrained baseline *A*_untrained_, representing a randomly initialized network where the self attention matrix is defined as

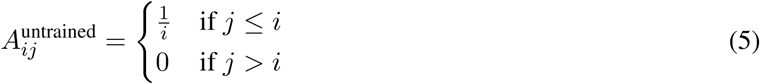

Using this baseline, we calculate the effective attention (*A*_eff_) as follows:

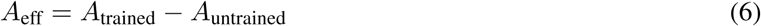

This normalization is an attempt to remove the architectural bias carried by the untrained model, so that what remains can be examined on its own.

The helicity proxy is computed from these normalized attention maps as follows. For every residue *i* we formed a profile by averaging *A* over both partners at each sequence separation,

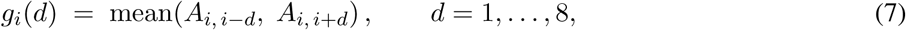

using whichever partners fall inside the sequence. Each profile was mean-centred, *g̃_i_*(*d*) = *g_i_*(*d*) – 〈*g_i_*〉*_d_*, so that what follows measures the shape of the profile rather than the overall level of attention at residue *i*. The window *d ≤* 8 is the smallest span covering two complete helical turns (7.2 residues). For each residue we ask how much of *g̃_i_*(*d*) is explained by a sinusoid at the *α*-helical frequency *ω* = 2*π/*3.6 rad per residue of separation. The Lomb–Scargle periodogram [29] fits, by least squares,

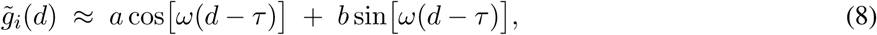

in which *a* and *b* are the fitting parameters and -the offset *τ* is defined by tan(2*ωτ*) = Σ_d_ sin(2*ωd*) Σ*_d_* cos(2*ωd*). The normalised power is the fraction of the profile’s variance captured by the fitted periodogram,

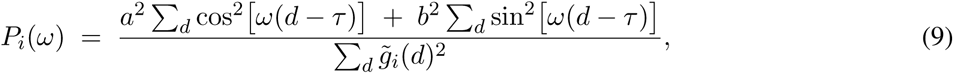

which is bounded in [0, 1] and equals the coefficient of determination *R*^2^ of the fit: *P_i_* = 1 when the profile is exactly a 3.6-residue wave, and *P_i_* = 0 when it carries no component at that frequency. The quantity we use as a computational proxy for helicity is the mean of this power over the peptide,

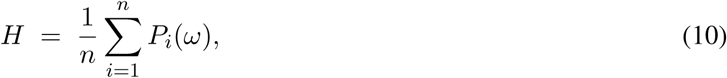

with *n* the number of residues. *H* is therefore the average degree to which a residue’s attention couples to sequence neighbours with the periodicity of an *α*-helical turn, and it is this single scalar per peptide that we correlate with the experimentally measured helicity.

These three techniques help us single out a cornerstone layer for our subsequent biological analysis. The extracted self-attention heatmaps help us identify amino acids with an important antimicrobial nature as well as correlations between various amino acids in the sequence. We tested our predictions on some well known and important antimicrobial peptides from the literature, and then applied the same analysis to the peptides generated by our own model.

### Data Sets

To train and evaluate our various models, we compiled datasets from previous literature and resolved inconsistencies, as much as possible. The sequence selection and MIC labeling rules are described below. Positive AMP sequences were obtained using identical collection pipelines from two distinct sources: Cordoves et al. [30] and the sequence dataset compiled in our prior work [31]. For sequences annotated with MIC values, a majority voting system was used to classify a peptide as antimicrobial. Specifically, if a peptide was associated with only a single MIC value across species, it was designated as an AMP if the MIC was less than 100 *µ*M. For peptides having multiple MIC values across different bacterial strains or species, we chose the peptide to be an AMP with a probability (*P*_AMP_):

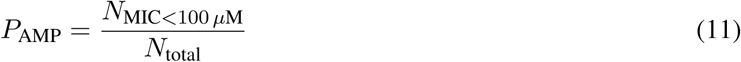

where *N*_MIC_*_<_*_100_ *_µ_*_M_ represents the number of MIC values below 100 *µ*M, and *N*_total_ is the total number of MIC values available for that peptide. Additionally, we collected peptides lacking explicit MIC data from the datasets provided in Ref. [30].

Table 1 summarizes the data collected for our various tasks. For training the generative model, we used only positive AMP sequences. For training the layer probing models, the probing classifier utilized positive AMPs and the non-antimicrobial peptide datasets compiled in Ref. [30], which were also used as the training data for our AMP-filtering classification models. The probing regressor was trained using datasets extracted from DBAASP V3 [32] as well as the dataset we collected previously [31]. Finally, the MIC regression tasks were trained on the exact dataset compiled in our previous manuscript [31]. Different datasets were used to match the information required by each task. Generative fine-tuning uses AMP sequences without additional labels, because the training targets are the amino acid tokens in the sequences themselves. In contrast, classifier training requires sequences labeled as AMPs or non-AMPs, while MIC regression requires sequences paired with measured MIC values.

**Table 1.** Comprehensive summary of dataset sources, model targets, and corresponding peptide selection criteria.

| **Model Framework** | **Specific Model / Purpose** | **Dataset Source** | **Peptide Selection Criteria** |
|---|---|---|---|
| Generative Modeling | Sequence Generation | Ref. [30] & Ref. [31] | (Positive) AMPs (MIC < 100 µM via majority vote) |
| Linear Probing | Probing Classifier<br>Probing Regressor | Ref. [30] & Ref. [31]<br>DBAASP V3 [32] | AMPs and Non-AMPs<br>Peptides without MIC values |
| AMP Filtering | Classification Models | Ref. [30] & Ref. [31] | AMPs, Non-AMPs |
| Probing Regression | Regression Models | Ref. [31] | Peptides with MIC values |

### Antibacterial Susceptibility Assays

The series of 18 peptides, that included 15 SI-generated peptides, the wild-type template (TP4) and two human-designed variants of it, were ordered with Biomatik (Kitchener, Canada) at a purity greater than 95%, in hydrochloride (HCl) salt form with less than 1% trifluoroacetate (TFA). Minimum inhibitory concentrations (MICs) of the generated peptides against the Gram-negative *Escherichia coli* MG1655 and the Gram-positive *Staphylococcus epidermidis* ATCC 14990 were determined using the broth microdilution assay as previously described [7]. Briefly, a fresh overnight bacterial culture was prepared. Peptides were serially diluted in a 96-well microtiter plate, followed by the addition of the bacterial suspension to achieve a final inoculum of 10^5^ CFU/mL. Wells containing only growth medium (without peptide or bacteria) served as negative controls. The plates were incubated at 37*^◦^*C for 24 h, after which bacterial growth was assessed by measuring the optical density at 600 nm (OD_600_) using a BioTek microplate reader (Agilent, Santa Clara, CA). Samples were prepared in triplicate and MICs were defined as the lowest peptide concentration that inhibited visible bacterial growth.

### Tryptophan Fluorescence Spectroscopy

Intrinsic tryptophan fluorescence of peptides was measured in both the absence and presence of lipid membranes in the form of small unilamellar vesicles (SUVs). The SUVs were composed of 1-palmitoyl-2-oleoyl-*sn*-glycero-3-phosphocholine (POPC) and the anionic 1-palmitoyl-2-oleoyl-*sn*-glycero-3-phosphoglycerol (POPG) at a 3:1 molar ratio and prepared as previously described [7]. 1,2-dimyristoyl-*rac*-glycero-3-methoxypolyethylene glycol-2000 (DMG-PEG_2000_) was included at 5% molar of total lipid to provide a steric barrier that prevents aggregation and fusion of the SUVs [33]. All peptide and lipid solutions were prepared in 5 mmol/L sodium phosphate buffer containing 5 mmol/L NaCl (pH 7.0). Lipid suspensions were serially diluted in a 96-well plate using an Opentrons OT-2 liquid handling system to generate a range of lipid concentrations. Peptides were added to each well at a final concentration of 10 *µ*M, resulting in lipid-to-peptide molar ratios (LPR) ranging from 3:1 to 200:1. An excitation wavelength of 280 nm was used, and emission was measured at 334 nm using a BioTek microplate reader with excitation and emission slit widths set to 10 nm and an optical gain of 100. Corrections for light scattering effects from SUVs were applied using a solution of tryptophan amino acid (Trp) suspended in the same SUV preparations, as described by Ladokhin et al [34]. Corrected fluorescence intensities at a fixed wavelength (334 nm) were normalized to the fluorescence intensity of the peptide in buffer and plotted against the lipid-to-peptide ratio (LPR) to produce binding curves (**Supplementary Fig S9**). The binding curves were fitted to a modified Langmuir binding model using a Markov Chain Monte Carlo sampling method to determine interaction parameters (Table 3) of the peptides with the lipid membranes, as described in detail elsewhere [35]. Results represent an average of two technical replicates (n=2).

### Circular Dichroism Spectroscopy

Secondary structures of the peptides in the presence and absence of lipid vesicles were analyzed using circular dichroism (CD) spectroscopy. Small unilamellar vesicles (SUVs) were prepared as described above and processed using an Opentrons OT-2 liquid handler in 96-well plates.

High-throughput CD measurements were performed using a Chirascan Q100 spectrometer (Applied Photophysics, Leatherhead, UK). Spectra were recorded across a wavelength range of 190–280 nm, using a 1 nm bandwidth and an integration time per point set to 2 s. The sample temperature was maintained at 25 *^◦^*C. Data were corrected by subtracting background contributions (SUVs in buffer alone), and the *α*-helical content of each peptide was calculated as previously described [7] (Table 3).

## Results

### Outcomes of the fine-tuning process

We initiated our pipeline by using LoRA to fine-tune the 1.2-billion-parameter ProGen model. To train the LoRA, we applied the LoRA weights on the Query and Key matrices of the model, bringing down the number of trainable parameters to less than 1%. Next we trained the model for 10 epochs across four NVIDIA A100 GPUs. The training time is approximately 30 mins for the model. Once trained, we sample sequences from the model. We utilized controlled top-p generation (*p* = 0.9) resulting in the generation of 20000 sequences in 10 minutes on a single A100 GPU.

To evaluate the impact of fine-tuning on generation quality, we generated 20,000 sequences from both the fine-tuned model and the baseline pretrained model using a maximum sequence length of 150. This sample size was chosen to match the scale of the fine-tuning training set (*N* 20,000). To assess distributional alignment, we evaluated the sequences generated by both models against the fine-tuning training set rather than a separate test set or the original base pretraining corpus. We selected the fine-tuning training dataset as the primary reference because its size provides a statistically robust, well-defined baseline representation of target physicochemical properties. By comparing the output distributions of the pretrained and fine-tuned models directly against this known training distribution, we measure how effectively fine-tuning shifts sequence generation toward the target domain. Once we generated these sequences, we calculated various physico-chemical properties in **Figure 2A** and show the amino acid compositions in **Figure 2B** and **S1**.

**Figure 2.**
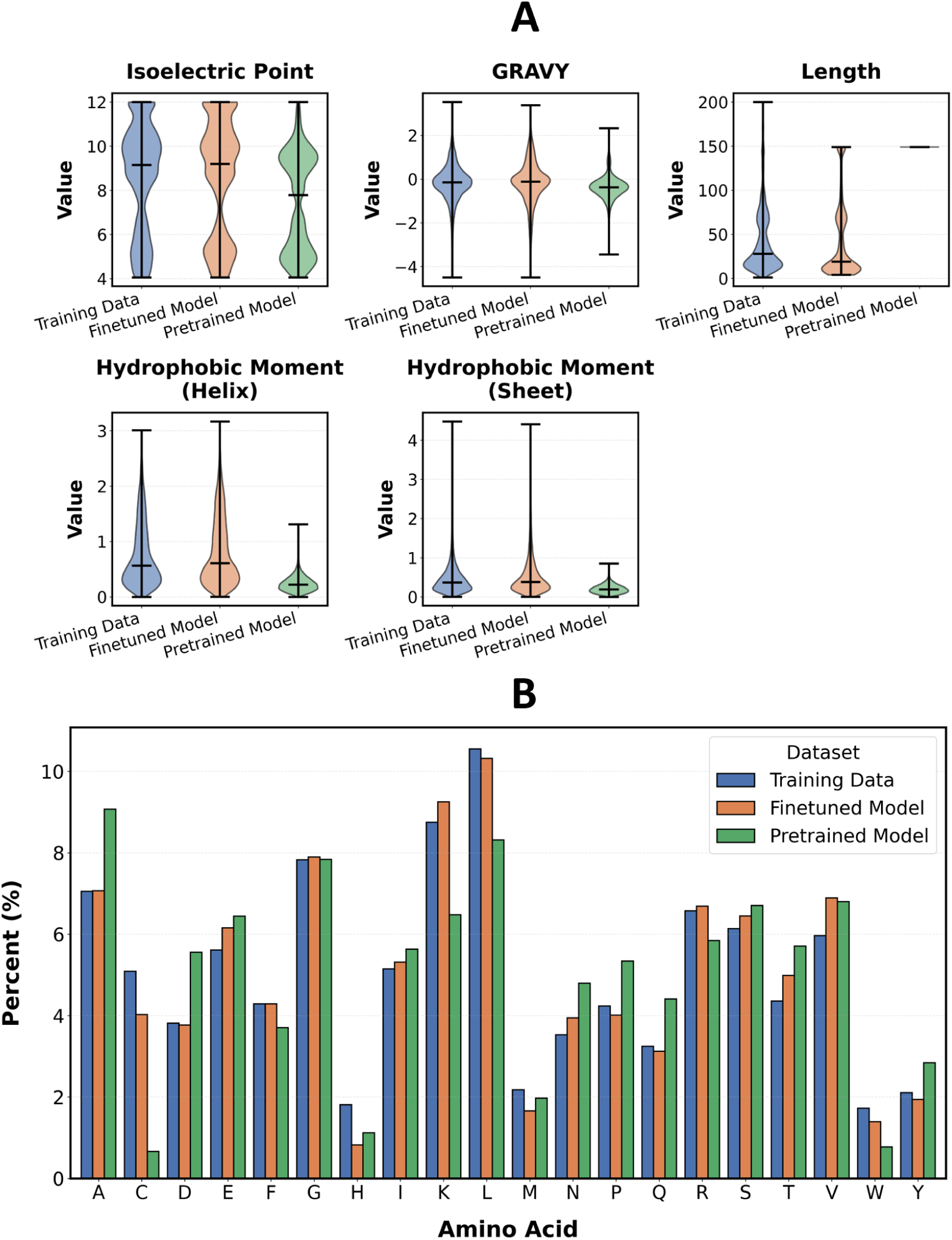
Comparative analysis of sequence fidelity: (A) Physico-chemical property distributions and (B) amino acid compositions for sequences derived from the training data (blue), fine-tuned model (orange), and pretrained model (green).

From **Figures 2A**, it is evident that the distribution of the physico-chemical properties of the sequences between the original training set and those generated through our fine-tuned model is much closer than the ones generated through the pretrained model. More specifically, there is a drastic discrepancy in the length of the sequences generated through our fine-tuned model compared to the baseline pretrained model. The pretrained model frequently attempts to generate sequences longer than 150 amino acids, resulting in outputs that are artificially truncated at the maximum seeded length of 150. Because the average length of natural proteins typically ranges from 300 to 500 amino acids, the baseline pretrained model naturally expects to generate much longer chains, causing it to frequently exceed our 150-amino-acid threshold. This further shows that fine-tuning indeed leads to improved generative capabilities of the model. The pretrained model learns from a broad protein corpus, with relatively limited AMP-specific examples, whereas our fine-tuning uses only AMP sequences. This adaptation shifts generation toward AMP-like lengths and sequence patterns.

Similarly, we looked at the compositions of the amino acids contained in the sequences obtained through the pretrained model, the training data and the fine-tuned model in **Figures 2A** and **S1**. While sequences generated through the pretrained model show more varied compositions as compared to the original AMP sequences (training data), the fine-tuned model better replicates the original training data. We observe that the differences between the original and fine-tuned sequences are particularly small for A, C, D, K, L, N, P, Q, and R.

### The setup for generation and filtering pipelines

Having established that fine-tuning shifts the generated sequence distribution towards that of natural AMPs, we used the model for de novo sequence generation. Candidate sequences were generated stochastically and subsequently routed through a multi-stage filtering pipeline. First, a binary classifier trained on ESM embeddings [30] was utilized to segregate true AMPs from non-antimicrobial sequences. The filtered, high-confidence AMP candidates were then evaluated using an ESM-based downstream regression model to predict their MIC values. For this analysis, we extracted ESM2 embeddings from AMP sequences and enhanced the base ESM2 model by adding two linear layers. A schematic of the SI model we developed to select candidate AMPs based on their predicted MIC is shown in **Figure 3**. Further details on the MIC prediction model and evaluation metrics are provided in prior work [36].

**Figure 3.**
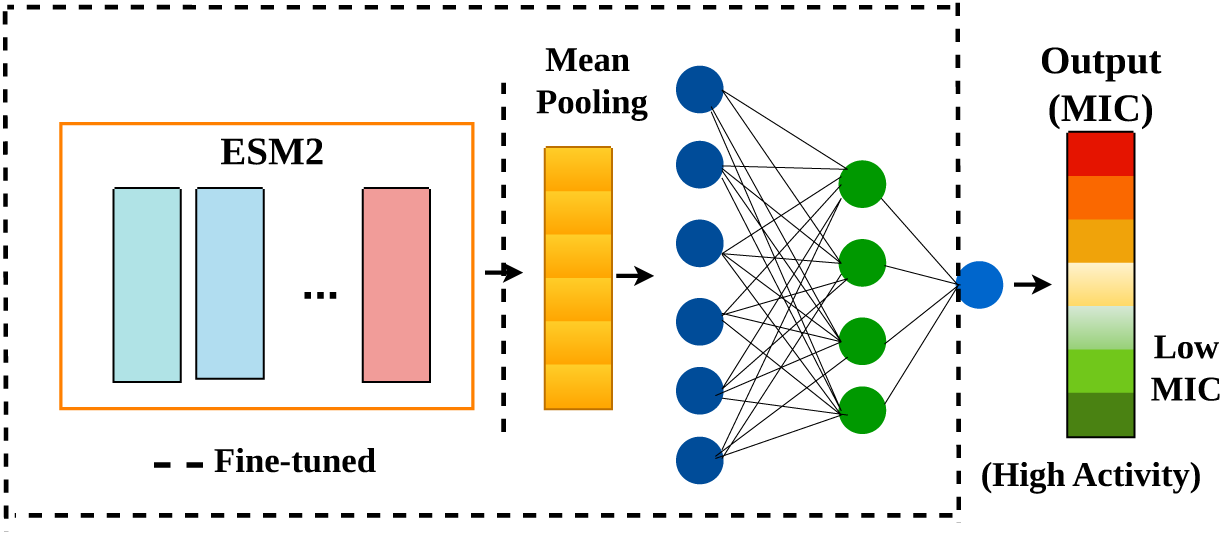
SI model used for MIC prediction analysis. We added two linear layers on top of ESM2 model and fine-tuned the entire model for MIC regression analysis.

### MIC and biophysical measurements

Following in silico MIC screening, we shortlisted 15 peptides for chemical synthesis and in vitro testing. We included three additional peptides based on structural rationale: the original wild-type TP4 peptide and two human-designed, slightly modified homologues of it. To facilitate controlled comparative analyses, the selection of candidates was constrained to sequences with ESM-derived numerical representations (embeddings) similar to that of Tilapia Piscidin 4 (TP4), a natural AMP that was previously characterized in our laboratory [7]. The synthesized variants were measured with biophysical and structural techniques that allowed quantification of their membrane interaction including experimental lipid-induced helicity and equilibrium binding affinities (*K_D_*). Finally, these empirical properties were mechanistically correlated with the model’s internal attention maps using our XSI framework to demonstrate the model’s ability in generating highly potent AMP sequences, and parse out a few key properties that define them.

Tables 2 and 3 summarize the measured peptides along with their biological activities in *E. coli* (MIC) and a few biophysical properties determined by CD and fluorescence spectroscopies (see Methods and Materials). The peptides are listed in the order of increasing MIC (decreasing antimicrobial potency). Notably, among the variants tested, eight of them (P1–P8) exhibit higher activity than the wild-type TP4 (P9), with MIC values lower than 4 *µ*mol/L. Thus, we confirm that the generative model, together with the filters we have designed, can successfully produce sequence homologues that show excellent antimicrobial potencies. Next, we wanted to uncover key physical and chemical attributes that make these homologues efficacious. For that, we examined their primary and secondary structural patterns, and quantifiable attributes that we can correlate with the biological activities. Because the lipid membranes of pathogens are the primary target for AMPs, we were interested in investigating these attributes in the presence of bacterial model membranes.

**Table 2.** Peptide sequence homologues to TP4, designed with the SI model. All peptides were SI-generated, with the exception of wild-type TP4 (P9), and two human-conceived variants (P10 and P18).

| **Name** | **Peptide sequence** |
|---|---|
| P1 | KWRSFIRGVTHAAHTALKHYFRRKRSG |
| P2 | LWGHIFRGIKHIGKTIHRLVTG |
| P3 | RWRSILKHGLKHIGKTIHRLVTG |
| P4 | VWRHFFRGIRHVGKTIHRLVTG |
| P5 | VWQHIFRGIVHVGRTIHRLITG |
| P6 | VTWRHIFRGIVHVGKTIHRLVTG |
| P7 | VKSWIRVAGRAIGKLIRRK |
| P8 | IWHHIVRGALHVGKLIHRLVTG |
| P9 | FIHHIIGGLFSAGKAIHRLIRRRRR |
| P10 | FIHHIIGGLFSAGRAIHRLIRRRRR |
| P11 | VTWRAIFRGIIHVGKTIHRLVTG |
| P12 | LWHIFKRIHRGIVHVGKTIHRLVTG |
| P13 | RWKGVIQAGKAIHRIIRRRR |
| P14 | VFWAKFKVAFRGVRHIGRTILRHVLR |
| P15 | VWGFIKHIVRPILRRLTRGLNKL |
| P16 | NWRNAWRRFKGAFRHVIHTAKRF |
| P17 | DWRFIRRIWKVGSHIYRAIKKLL |
| P18 | FIHHIIWILFSAGRAIHRLIRRRRR |

We first measured the binding characteristics of the peptides by tryptophan fluorescence spectroscopy in the absence and presence of lipid SUVs in buffer. With the exception of P9 and P10, all peptide variants contained W in their sequences. Upon interacting with lipid bilayers, the emission spectrum of W displays an increase in fluorescence intensity, as explained in detail by Ladokhin et al. [34] (see Methods). Serial dilutions of lipid SUVs were prepared and incubated with the individual W-containing peptides. The W fluorescence intensity was measured at a fixed excitation and emission wavelength and plotted as a function of lipid concentration to generate binding curves (**Figure S9**). Past a certain lipid to peptide ratio (LPR), saturation is reached and the intensity plateaus at *I*_max_ which is, broadly speaking, associated with the amount of peptide bound at equilibrium, in excess lipid ( **S9**). The binding curves also contain information about the number of lipid headgroups bound per peptide (n) and the dissociation constant *k_D_*, which can be, to best approximation, derived from fitting a binding model to the curves (Methods, and **Figure S9**).

Further, to evaluate membrane-induced changes in peptide secondary structure, we performed CD spectroscopy, for peptides in the absence of SUVs (in buffer) and with SUVs (in lipid), at an LPR of 1/100, where it appears that all peptide binding events have reached saturation (**Figure S9**). As it is the case for many AMPs, TP4 transitions from an unstructured peptide in buffer to a partial, and in some cases, fully helical peptide in the presence of lipid bilayers [7], [35]. As shown in Table 3, all 18 peptides show low helical content (i.e., the fraction of a peptide sequence that folds into a helix) in buffer, but exhibit a significant increase in helicity upon interacting with the lipid membrane, over a wide range of values (Table 3).

Given the spectrum of measured biophysical parameters that we are using here to report on peptide-membrane interactions, and biological activities (MIC values) displayed by the 18 peptides, we resorted to calculating a correlation matrix between these parameters (**Figure 4**).

**Figure 4.**
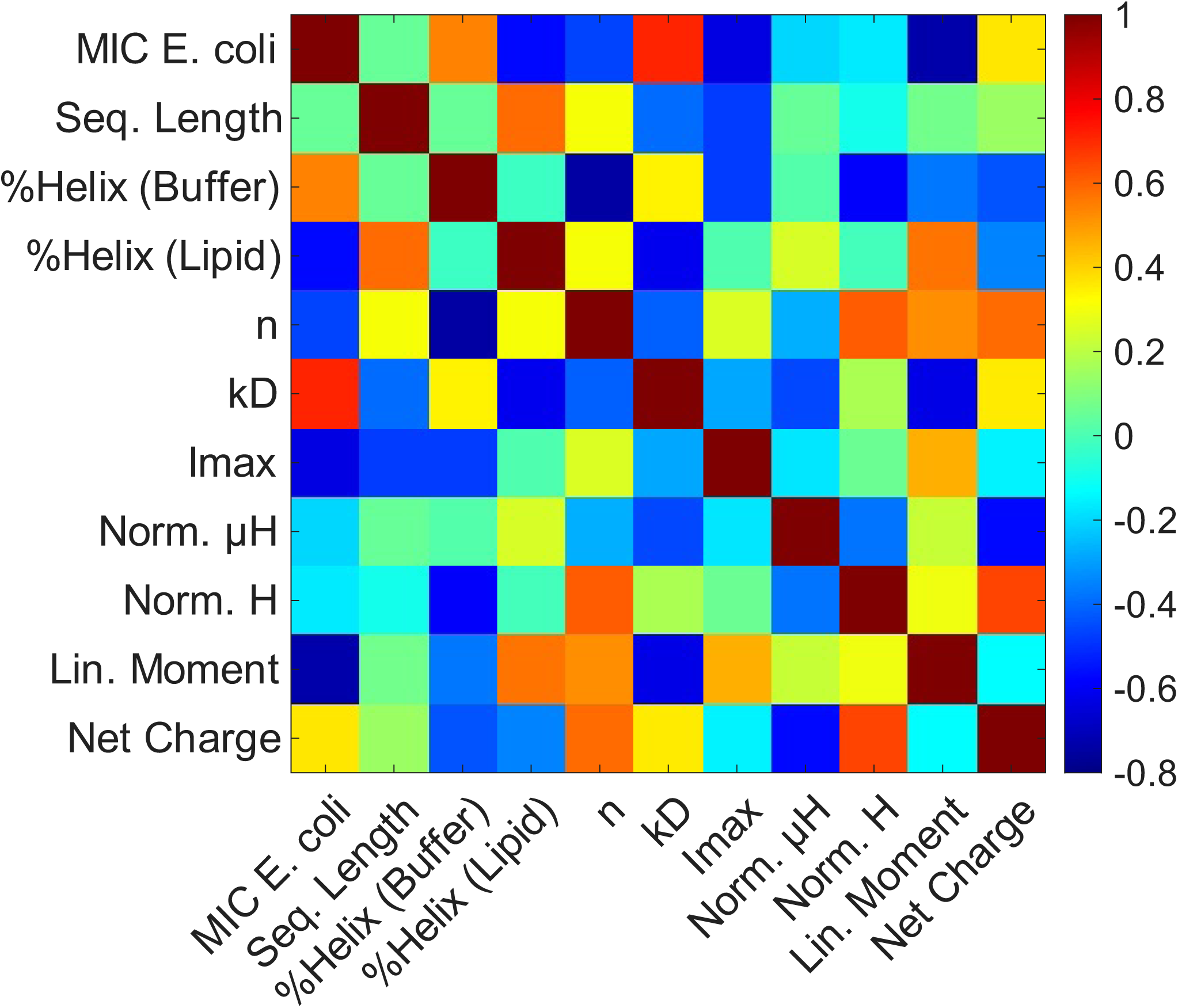
A correlation map between measured and calculated, biophysical parameters for the SI-generated TP4 peptide homologues. Intense/dark red indicates strong, positive correlation, such as that observed between the MIC and the dissociation constant, *k_D_*(lower MIC equate with lower *k_D_*). Blue indicates negative correlations (lower MIC equates with greater helical content of the peptide in the lipid environment). A few properties, calculated from the sequences, such as peptide charge, sequence length and helical and linear hydrophobic moments, are also included. They were calculated using the DBAASP tool (Ref [32]). The parameter values are shown in Table 3.

**Table 3.**
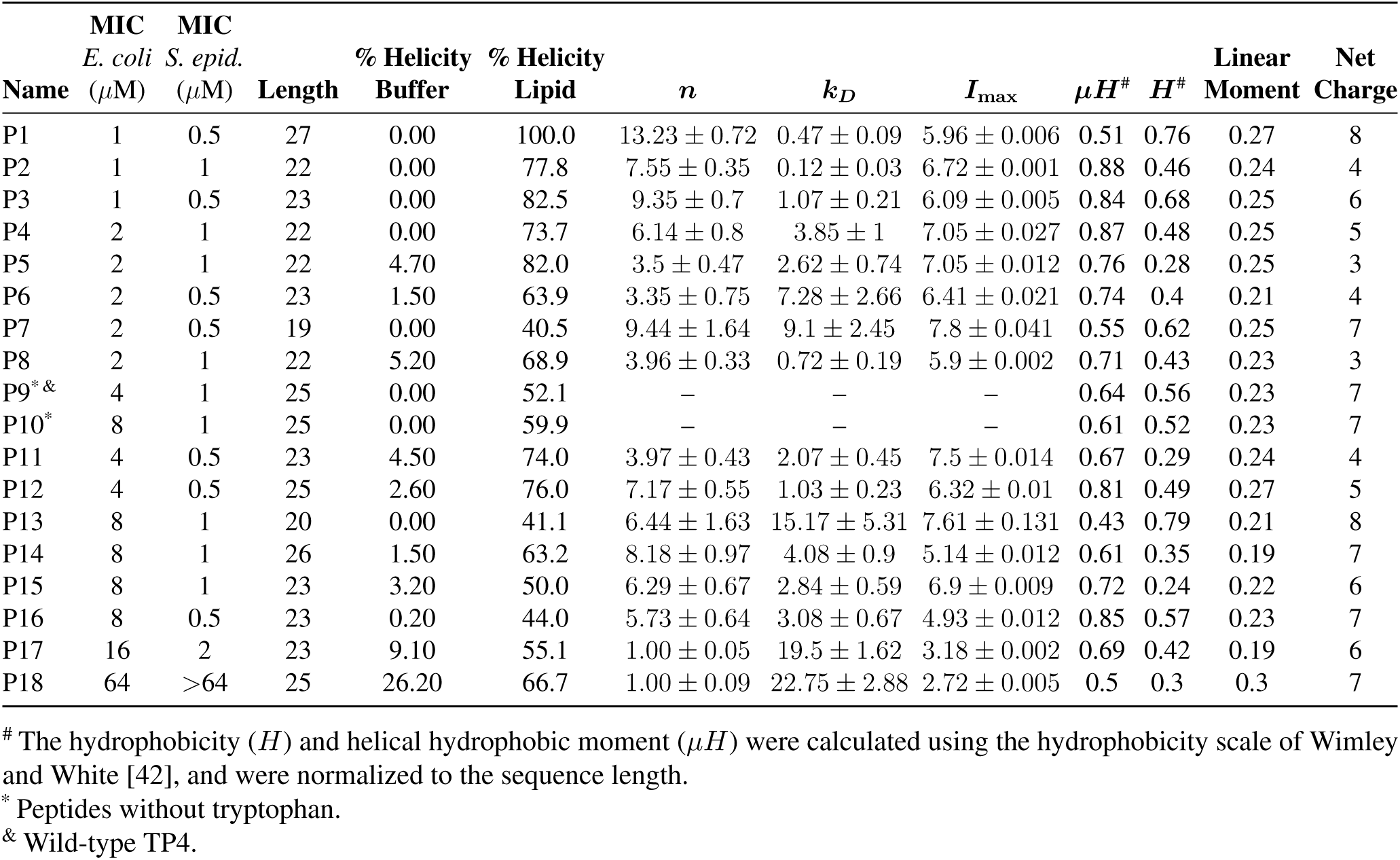
Biological and biophysical parameters for the investigated peptide variants. Fit parameters are defined as: *n* = number of lipids bound per peptide; *k_D_* = dissociation constant; *I*_max_ = the maximum fluorescence intensity, reached at saturation.

| **Name** | **MIC**<br>*E. coli*<br>(µM) | **MIC**<br>*S. epid.*<br>(µM) | **Length** | **% Helicity**<br>**Buffer** | **% Helicity**<br>**Lipid** | ***n*** | ***k<sub>D</sub>*** | ***I***<sub>max</sub> | ***µH***<sup>#</sup> | ***H***<sup>#</sup> | **Linear**<br>**Moment** | **Net**<br>**Charge** |
|---|---|---|---|---|---|---|---|---|---|---|---|---|
| P1 | 1 | 0.5 | 27 | 0.00 | 100.0 | 13.23 ± 0.72 | 0.47 ± 0.09 | 5.96 ± 0.006 | 0.51 | 0.76 | 0.27 | 8 |
| P2 | 1 | 1 | 22 | 0.00 | 77.8 | 7.55 ± 0.35 | 0.12 ± 0.03 | 6.72 ± 0.001 | 0.88 | 0.46 | 0.24 | 4 |
| P3 | 1 | 0.5 | 23 | 0.00 | 82.5 | 9.35 ± 0.7 | 1.07 ± 0.21 | 6.09 ± 0.005 | 0.84 | 0.68 | 0.25 | 6 |
| P4 | 2 | 1 | 22 | 0.00 | 73.7 | 6.14 ± 0.8 | 3.85 ± 1 | 7.05 ± 0.027 | 0.87 | 0.48 | 0.25 | 5 |
| P5 | 2 | 1 | 22 | 4.70 | 82.0 | 3.5 ± 0.47 | 2.62 ± 0.74 | 7.05 ± 0.012 | 0.76 | 0.28 | 0.25 | 3 |
| P6 | 2 | 0.5 | 23 | 1.50 | 63.9 | 3.35 ± 0.75 | 7.28 ± 2.66 | 6.41 ± 0.021 | 0.74 | 0.4 | 0.21 | 4 |
| P7 | 2 | 0.5 | 19 | 0.00 | 40.5 | 9.44 ± 1.64 | 9.1 ± 2.45 | 7.8 ± 0.041 | 0.55 | 0.62 | 0.25 | 7 |
| P8 | 2 | 1 | 22 | 5.20 | 68.9 | 3.96 ± 0.33 | 0.72 ± 0.19 | 5.9 ± 0.002 | 0.71 | 0.43 | 0.23 | 3 |
| P9<sup>*&</sup> | 4 | 1 | 25 | 0.00 | 52.1 | – | – | – | 0.64 | 0.56 | 0.23 | 7 |
| P10<sup>*</sup> | 8 | 1 | 25 | 0.00 | 59.9 | – | – | – | 0.61 | 0.52 | 0.23 | 7 |
| P11 | 4 | 0.5 | 23 | 4.50 | 74.0 | 3.97 ± 0.43 | 2.07 ± 0.45 | 7.5 ± 0.014 | 0.67 | 0.29 | 0.24 | 4 |
| P12 | 4 | 0.5 | 25 | 2.60 | 76.0 | 7.17 ± 0.55 | 1.03 ± 0.23 | 6.32 ± 0.01 | 0.81 | 0.49 | 0.27 | 5 |
| P13 | 8 | 1 | 20 | 0.00 | 41.1 | 6.44 ± 1.63 | 15.17 ± 5.31 | 7.61 ± 0.131 | 0.43 | 0.79 | 0.21 | 8 |
| P14 | 8 | 1 | 26 | 1.50 | 63.2 | 8.18 ± 0.97 | 4.08 ± 0.9 | 5.14 ± 0.012 | 0.61 | 0.35 | 0.19 | 7 |
| P15 | 8 | 1 | 23 | 3.20 | 50.0 | 6.29 ± 0.67 | 2.84 ± 0.59 | 6.9 ± 0.009 | 0.72 | 0.24 | 0.22 | 6 |
| P16 | 8 | 0.5 | 23 | 0.20 | 44.0 | 5.73 ± 0.64 | 3.08 ± 0.67 | 4.93 ± 0.012 | 0.85 | 0.57 | 0.23 | 7 |
| P17 | 16 | 2 | 23 | 9.10 | 55.1 | 1.00 ± 0.05 | 19.5 ± 1.62 | 3.18 ± 0.002 | 0.69 | 0.42 | 0.19 | 6 |
| P18 | 64 | >64 | 25 | 26.20 | 66.7 | 1.00 ± 0.09 | 22.75 ± 2.88 | 2.72 ± 0.005 | 0.5 | 0.3 | 0.3 | 7 |

<sup>#</sup> The hydrophobicity (*H*) and helical hydrophobic moment (*µH*) were calculated using the hydrophobicity scale of Wimley and White [42], and were normalized to the sequence length.
<sup>\*</sup> Peptides without tryptophan.
<sup>&</sup> Wild-type TP4.

In addition to the experimental measurements, we reported sequence-derived attributes relevant to AMP–membrane interactions [37]. These parameters include: the peptide charge - important for recruiting the cationic AMP to the negatively charged bacterial membranes; the hydrophobic moment (*µ*H) – a measure of the amphipathicity due to clustering of polar residues on one side of the helix and hydrophobic residues on the opposite side, thereby, facilitating insertion into the bilayer [38]; the linear hydrophobic moment [39] – a measure of the separation of polar and hydrophobic amino acids along the primary sequence of the peptide chain (e.g., a completely symmetrical peptide sequence has a linear moment equal to zero).

Although XAMPGen has focused more on sequences active against *E. coli*, which was due to the higher availability of training data for this bacterial species, we also measured the antibacterial activities against the Gram-positive *S. epidermidis*, for comparison (Table 3). We see that many of the generated peptides are highly active against both bacterial species, although their correlations with the measured parameters differ. These results imply that differences in the membrane composition between Gram-negative and Gram-positive bacteria may influence AMP–membrane interactions and their mechanisms of action.

Strong, positive correlations are seen between the antimicrobial potency and their binding affinities to the membranes (e.g., the lower the dissociation constant *k_D_*, the lower the MIC). Strong, negative correlations are observed between the antimicrobial potency and both the amount of lipids bound per peptide, and the peptide helicity in the presence of lipid (e.g., the greater the *n*, *I*_max_ or percent helicity, the lower the MIC). Interestingly, the helical hydrophobic moment displays a weak correlation with the antibacterial activity. This may be explained by the fact that the calculations of *µ*H are usually inaccurate without knowledge of the exact 3D structure of the peptides, and rely heavily on extrapolations from a very limited set of available 3D structures of AMPs in bilayers. We included it here, as it is often cited as one of the key drivers of peptide insertion, and membrane perturbation, leading to lysis [40, 41]. The linear hydrophobic moment, which does not presuppose a helical conformation and is therefore the more reliably calculated of the two, tracks the activity considerably better, and is the sequence-derived quantity most strongly related to the MIC in **Figure 4**.

This observation aligns with the clustering of R residues (polyR) toward the end of the sequence, discussed below in our XSI analysis, that we have shown to be a remarkable feature displayed by TP4 and several other natural peptides. It plays key roles in damaging the bacterial membranes through mechanisms such as domain formation and LPS sequestration in bacterial membrane mimics [7]. By starting from the TP4 model and working in a closed space around this template, XAMPGen appears to have captured the separation of (hydrophobic – charged) amino acids along the chain, as important for enhanced action at bacterial membranes. Taken together, our measured data and analyses support a common mechanism governing the activity of the TP4-like peptides: peptides exhibiting greater activity are those that bind bacterial membranes most strongly and undergo the largest membrane-induced conformation transition. Although the SI-generated peptide set is limited, it does point to key sequence properties that, when present, enhance the antibacterial efficacy. A few of these important attributes, revealed through the attention heatmaps, may be difficult to observe otherwise and warrant more experimental exploration.

### XSI analysis

#### Layer probing

Having generated, synthesized and experimentally characterized the peptides, we now turn to the model itself, to gain mechanistic insight into its internal representations. By using techniques from XSI commonly used to study LLM, we dove deeper into the information flow within these networks using layer probing, an approach that has been used quite recently to study LLMs in the NLP domain [43, 44, 45, 46]. To probe ProGen’s hidden layers, we used three probing tasks: (i) a linear probing classifier to distinguish AMP vs non-AMP, (ii) a linear probing regressor to predict the MIC values of the peptides, and (iii) a structural probe that compares the helicity read off a layer’s attention map with the helicity measured in membranes by CD spectroscopy. For the first two probing tasks, we pass sequences through the model and then fit some linear layers on top of the extracted hidden representations to measure the performance of each layer towards the downstream task. To evaluate layerwise representations, we employ four standard metrics. For binary classification—designating AMPs as the positive class (1) and non-AMPs as negative (0)—performance is measured using Accuracy [47] and Matthews Correlation Coefficient [48] (MCC), derived from True Positives (*TP*, correctly identified AMPs), True Negatives (*TN*, correctly identified non-AMPs), False Positives (*FP*, non-AMPs predicted as AMPs), and False Negatives (*FN*, AMPs predicted as non-AMPs):

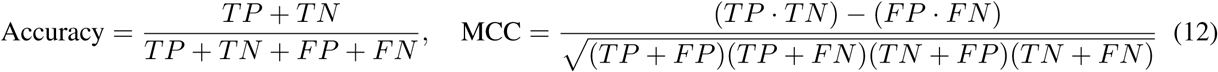

While Accuracy reflects overall correctness, MCC accounts for class imbalance across all confusion matrix categories (TP, TN, FP and FN). For MIC regressor probing, performance is evaluated using the Coefficient of Determination (*R*^2^) [49] and Pearson Correlation Coefficient (PCC) [50]:

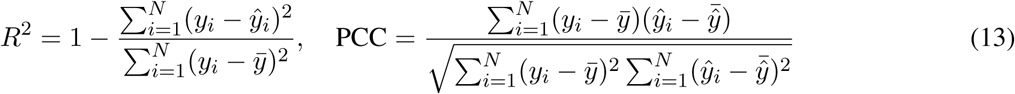

where *y_i_*and *ŷ_i_* denote ground-truth and predicted MIC values for sequence *i*, with sample means *y̅* and 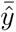. *R*^2^ quantifies explained variance, while PCC assesses linear alignment. For the third probing task, we use the normalized attention maps obtained from each layer: the attention heatmaps are averaged over all heads and normalized as in Equations 5 and 6, a helicity is computed for every peptide from every layer through Equations 7–10, and these values are compared with the helicities measured experimentally, the agreement being quantified by the PCC.

In **Figure 5A**, we show the layer performance for the classification task of the linear probes, while in **Figures 5B** and **5C** we show the layer performance for the regression probe. Across these two tasks, we observed a distinct “U-shaped” performance trajectory. Initial layers exhibit high predictive power, followed by a noticeable performance drop in the intermediate layers. The performance subsequently recovers, peaking steadily in the final layers (above layer 11). This trajectory reflects the inherent architecture of deep transformers. While the early layers directly process the initial input embeddings, the intermediate layers lack a direct interface with either the inputs or the final output objective, causing linear separability to temporarily drop. Subsequently, the later layers consolidate these complex abstractions into the context-rich representations necessary to finalize sequence generation, rendering them highly performant for our downstream probes.

**Figure 5.**
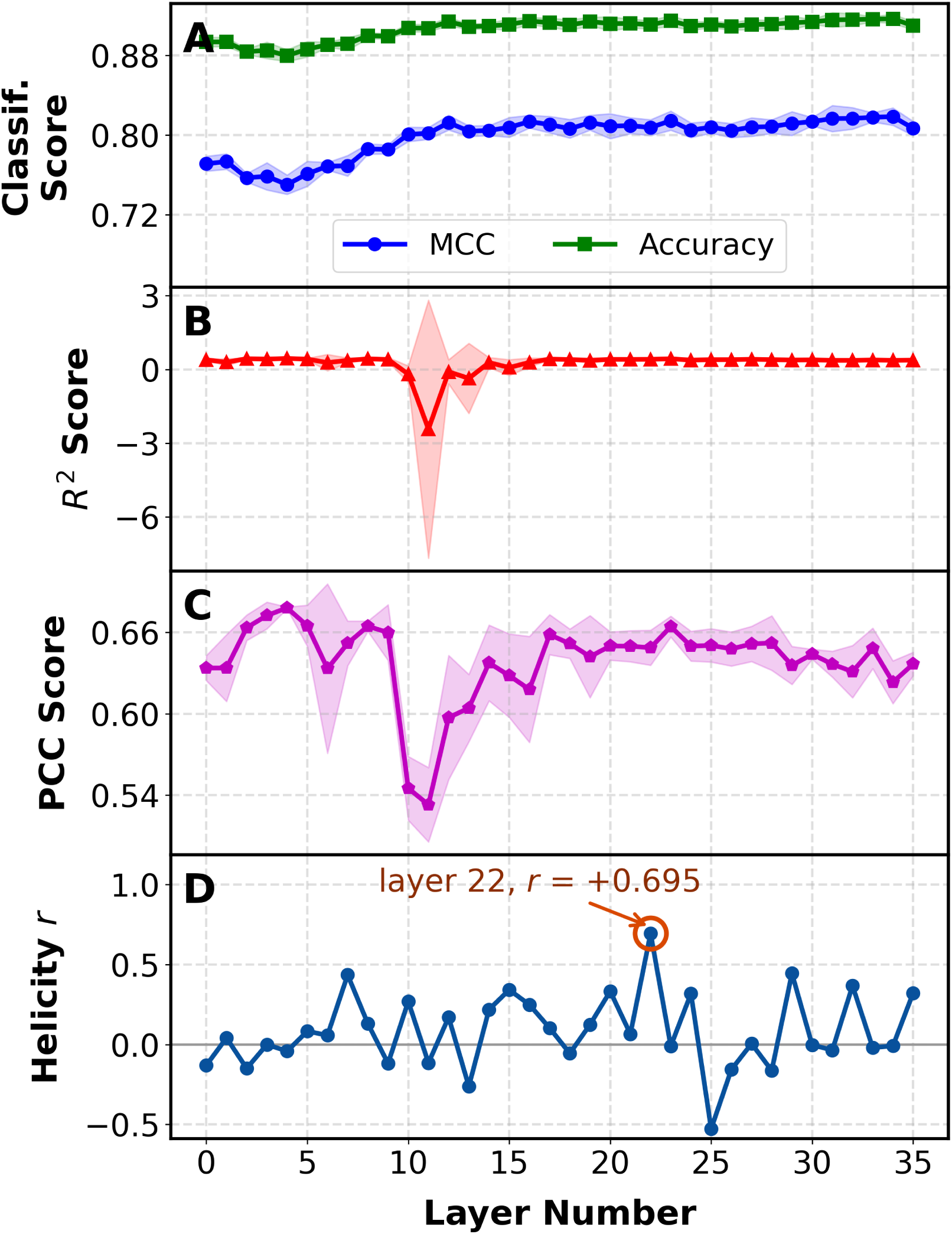
The three probing tasks, at every one of the 36 layers. (A) linear classification probing; (B) and (C) linear regression probing for MIC, scored by *R*^2^ and by the PCC. Scores in (A)–(C) are means over five probe initialisations, the bands giving the seed-to-seed standard deviation. (D) the helicity probe: the Pearson correlation, across the 18 measured peptides, between the attention helicity proxy of Equation 10 and the CD helicity in lipid of Table 3. The first two probes plateau over the later layers without singling one out; the helicity probe peaks sharply at layer 22 (circled), *r* = +0.695 (*p* = 0.0014), no other layer reaching *r* = 0.444.

In both the classification and regression tasks, later layers perform consistently well, making it challenging to select a cornerstone layer for detailed biological analysis. We therefore used a different probe, the helicity probe, whose results are shown in **Figure 5D**. This probe revealed larger discrepancies between layers, with layer 22 exhibiting the strongest correlation with experimentally measured helicity. Since our manuscript mainly focuses on AMPs that form helices on interacting with membranes, we selected layer 22 for subsequent analysis. However, in the section on the more granular approach, we show that layer 22 alone cannot capture all of the relevant biological insights, and that looking at specific heads and layers can single out specific biology.

Layer 22 shows a Pearson correlation of *r* = 0.695 (*p* = 0.0014) and a Spearman correlation of *ρ* = 0.738 (*p* = 0.0005) with the measured helicities across the 18 peptides. In **Figure 6**, we show the correlation between the helicity values, experimental and computational, for all the peptides measured through CD.

**Figure 6.**
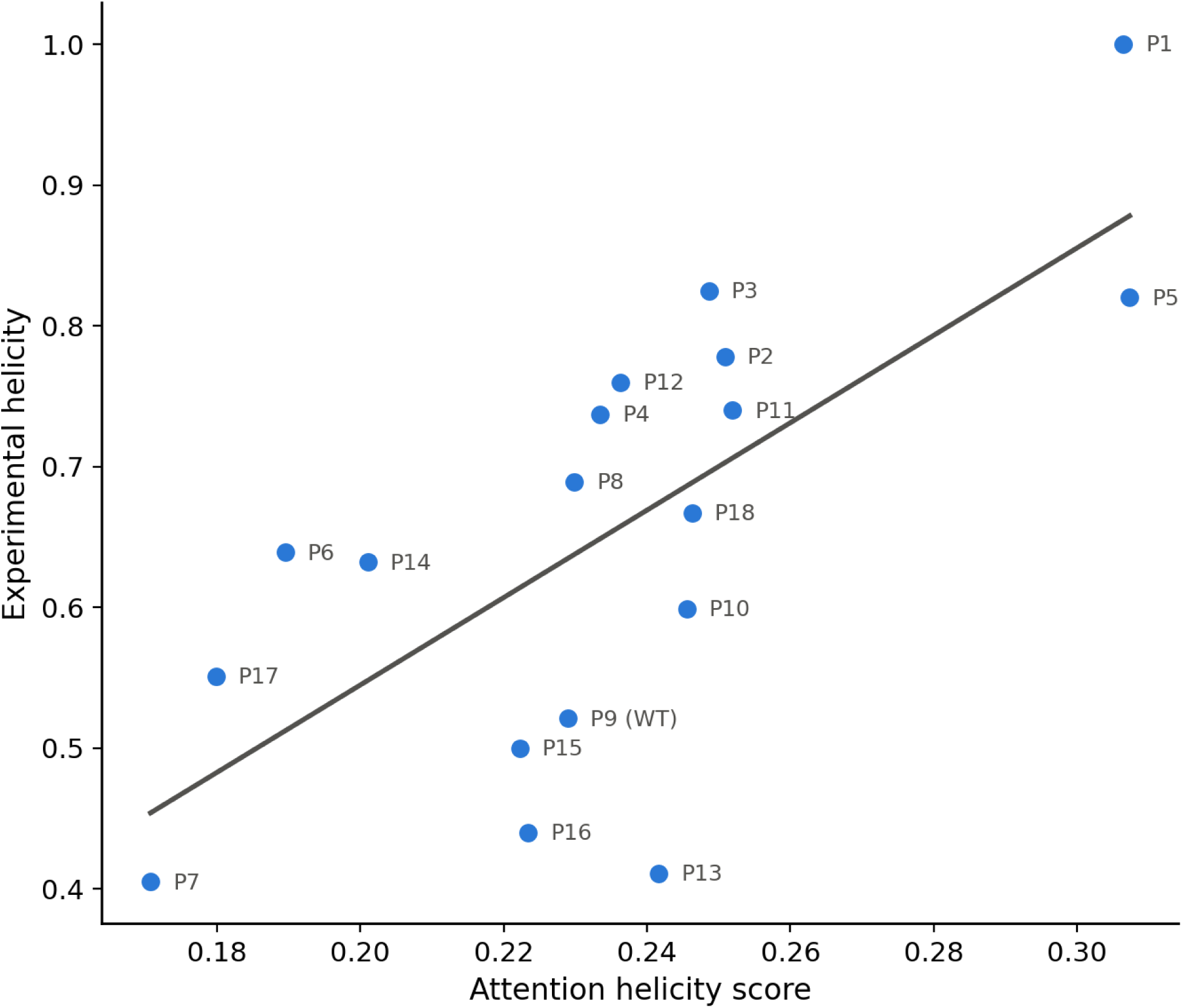
The correlation between experimental and computed, corrected helicities.

As a comparison of our peptide helicity proxy against standard structural predictors, we used two freely available tools from the literature, ESMFold [15] and OmegaFold [51], folding the same 18 sequences and assigning the helix fraction of every predicted structure with DSSP [52]. We outperform both methods: against the CD helicities ESMFold reaches only *r* = +0.268 (*p* = 0.28) and OmegaFold *r* = +0.216 (*p* = 0.39). Each returns an almost uniformly helical model for every peptide — helix fractions of 0.89 0.06 for ESMFold and 0.89 0.02 for OmegaFold against a measured spread of 0.41–1.00 — so neither model accurately predicts helical content. This is likely because ESMFold and OmegaFold are trained on proteins, and more generally on soluble proteins, which are not necessarily membrane-active. Layer 22 is therefore adopted as the cornerstone layer for all attention analyses that follow.

Once we have probed the LLM to identify the layers containing the highest concentration of information, we turn to the self-attention maps of the model to examine the biology it may have learnt. Because the ProGen model is large and deep, inspecting all of its self-attention maps is impractical. To mitigate this issue, we adopted two strategies for analyzing the most critical attention maps: (i) averaging across all attention heads, and (ii) restricting the analysis to layer 22.

### Attention heatmap analysis on known antimicrobial peptides

As in natural language models, the amino acids in a peptide sequence can be treated like the words in a sentence: both identity and position carry meaning, and how much a residue matters depends on its context. ProGen is a deep, decoder-only transformer of 36 stacked self-attention layers, trained on next-token likelihood (Eq. 1); it never scores residues itself, so any notion of importance is one we read out post hoc from its attention. We do so at layer 22 as shortlisted from the previous Layer Probing section, averaging over all attention heads, and illustrate the result for several well-characterized AMPs (**Figure 7**).

**Figure 7.**
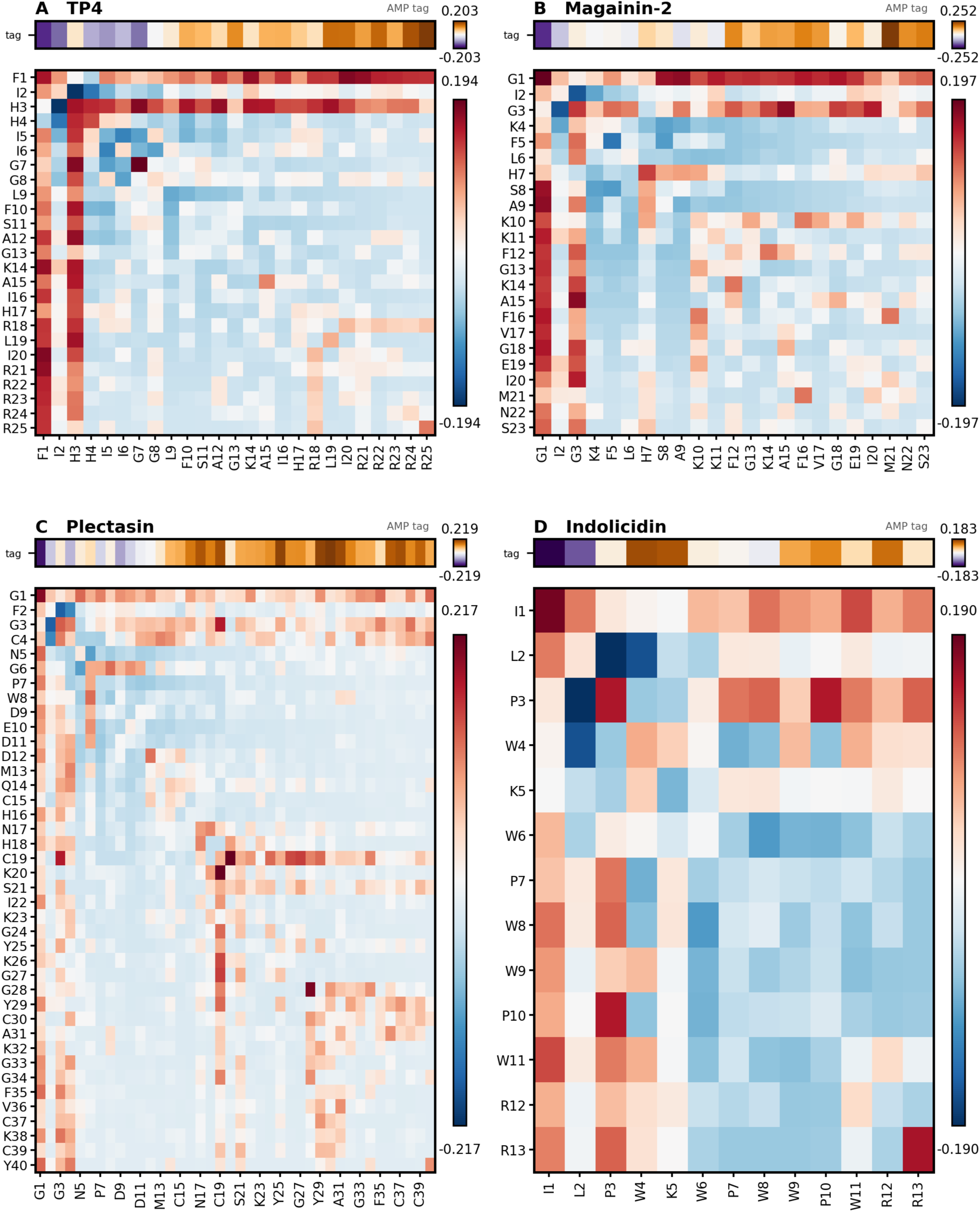
A visualization of the effective attention matrix at layer 22, describing the amino acid interdependencies within a peptide sequence and the importance of each amino acid for the “AMP” tag, for: (A) Tilapia piscidin-4 (TP4), FIHHIIGGLFSAGKAIHRLIRRRRR; (B) Magainin-2, GIGKFLHSAKKFGKAFVGEIMNS; (C) Plectasin, GFGCNGPWDEDDMQCHNHCKSIKGYKGGYCAKGGFVCKCY; (D) Indolicidin, ILPWKWPWWPWRR.

Each map shows how much attention the model pays to each residue while processing every other residue. Because ProGen reads a sequence in one direction only, residues near the N-terminus spread their attention over fewer preceding tokens and would otherwise appear artificially prominent; we therefore subtract the attention expected by chance (Eqs. 5 and 6), so that each value reports whether the model attends to a residue more or less than an even distribution would predict. In **Figure 7**, red colored cells in the attention maps denote higher attention than chance, blue denotes lower attention than chance, and white, near chance. The color-bar values are unitless differences in attention weights, and they have meaning only within the individual panel: each panel carries two of them, one for the residue–residue map and one for the tag strip above it, and neither transfers to another peptide. Rows and columns of the map list residues in sequence order, labelled by amino acid and position (e.g., R21 represents the arginine in position 21 of the TP4 sequence, **Figure 7A**). The narrow strip above each map, labelled “tag”, gives each residue’s attention to the antimicrobial control tag, which we use here to point to its association with the antimicrobial character. Every cell in the map gives the coupling between two residues at their specific positions, so that a red-colored cell at row R21, column F1 means that the model recognizes the relative importance of having a phenylalanine in position 1 (F1) and an arginine in position 21 (R21). Because each row sums to zero after correction, the diagonal is fully determined by the rest of its row and is not interpreted. These patterns are statistical regularities learned from sequence alone: they show where the model attends, and serve here as hypotheses to be tested experimentally.

We start our attention map analysis by focusing on the TP4 peptide, an important AMP that was characterized in detail, in our laboratory (**Figure 7A**). TP4 is one of the most potent members of the piscidin family of AMPs that are widely spread in teleost fish ( [53]). The piscidin family (Pfam 08107) comprises 413 catalogued proteins in UniProt [54]. Using a local BLAST+ search [55], we scanned their mature peptide regions (signal peptides removed) against our training set and found 290 piscidin like proteins (60% coverage *E <* 0.20). They are characterized by high sequence conservation that includes a high content of histidine (H) residues, a N-terminal Cu^2+^/Ni^2+^ binding motif (ATCUN motif) which requires a histidine in position 3, and a series of concatenated arginine residues (polyR) at the C-terminal tail. We first turn our attention to this polyR feature. In our previous experimental work [7], we demonstrated that this polyR tail (R21–R25) is crucial for antimicrobial activity due to the strong electrostatic affinity of this motif for negatively charged bacterial membranes. This important motif, frequently found in piscidins, and its biological manifestation is clearly reflected in our attention heatmap analysis (**Figure 7A**), which highlights the value brought by this segment alone to an AMP. The importance of the polyR segment is unmistakable in the heat map analysis and is further suggested by applying virtual mutations in our model. For instance, replacing the polyR tail with a poly-lysine (polyK) causes a single lysine residue (K22) to become highly dominant, while simultaneously disrupting the correlation between the preceding R18 residue and the R21–R25 region (**Figure S2A**). Overall, however, the substitution changes the map very little, all differences in effective attention between the two heatmaps staying below 0.09, as shown by the difference map of **Figure S3**. Conversely, truncating the polyR tail entirely does not induce major changes in the global attention matrix of the remaining core sequence (**Figure S2B**). This indicates that the polyR tail acts as an independent functional entity, confirming our previous findings for this peptide family [7].

Moving our attention from arginines to histidines, we see that for all three piscidin variants, the heatmaps (**Figures 7A**, **S2A**, and **S2B**) display a distinct vertical dark band persists at the H3 column, indicating strong sequence-wide correlations. While H3 does not directly dictate antimicrobial efficacy, it belongs to the ATCUN motif that is biologically important for the AMP’s mechanisms of action, by carrying metal ions inside cell, destabilizing membranes and generating reactive oxygen species [56]. Apart from that we also find that the model assigns high importance to residues like H17 which is highly conserved in Piscidin peptides and it might lead to higher permealization in piscidin peptides [57]. Similar to H3 and H17, the N-terminal phenylalanine-1 (F1) residue displays broad correlation with the rest of the chain. From a biophysical perspective, highly hydrophobic and aromatic residues such as F or W, positioned toward the N-terminus, are known for their role in membrane insertion, anchoring the peptides in the hydrocarbon region of the lipid membrane, and participating in amphipathic, helical structure formation of AMPs at the bilayer-water interface [58], [59].

Next, we examined Magainin-2 (**Figure 7B**). The N-terminal glycine residues (G1 and G3) emerge as important amino acids, each appearing as a broad band of positive coupling across the sequence, possibly due to the model capturing their role in conferring flexibility and initiating the N-terminus binding to the membrane, together with I2 [60] [61]. This ‘GIG’ motif at the start of the sequence appears also in the archetypal AMP, Melittin. Furthermore, as for most helical AMPs, a balance of hydrophobic and polar amino acids have to exist that allow the peptide to fold into an amphipathic helix, characterized by a hydrophobic moment. The hydrophobic moment of Magainin-2 has been related to the presence of both hydrophobic and hydrophilic amino acids, which explains why both types are dominant in the attention matrices (S, N, K, and E for polar and F, A, V, I, and M for hydrophobes) [62].

For the defensin Plectasin (**Figure 7C**), the attention map highlights residues F2, G3, C4 and C37, which hydrogen-bond the pyrophosphate moiety of the bacterial cell-wall precursor Lipid II — the target through which plectasin acts [63]. Interestingly, plectasin has six cysteines forming three covalent disulfide bridges (Cys4–Cys30, Cys15–Cys37, and Cys19–Cys39) [64] that endow the peptide with remarkable stability against enzymatic degradation and high bioactivity against bacteria. The matrix further registers positions concentrated in the helix C-terminus and the *αβ*-loop (H16, H18, K20, I22, K23, Y25, K26). Recent experiments shows that this loop is central to the mechanism: I22 and Y25 form a clamp that locks it in a primed conformation, and the mutants I22A and Y25K abolish Lipid II binding entirely [65]. Two of Plectasin’s three disulfide bonds (C4–C30 and C19–C39) are also mapped in our matrices.

We also analyzed the indolicidin (**Figure 7D**), a small, 13-amino acid antimicrobial peptide found in cow neutrophil blood cells. The peptide is particularly rich in tryptophan and proline, and also displays two R’s at the C-terminus. In contrast to the majority of AMPs, indolicidin remains largely unstructured in interaction with membranes, and while it can cross membranes, does not lyse them. Just like F, W is a large hydrophobic residue known to help stabilize peptide conformation and anchor AMPs to target cell membranes, with a strong preference for the bilayer-water interface [66], which show up in our attention maps as highly correlated to the other residues across the sequence. Furthermore, W and R amino acids can work together to target and lock AMPs in membranes via cation-*π* interactions [67]. On the other hand, Proline is known for conferring peptide flexibility by preventing formation of rigid *α*-helices [68]. Proline is found in AMPs that kill bacteria via non-lytic mechanisms [69]. The strong W-P correlations in the heatmap are consistent with these structural roles, although indolicidin’s high tryptophan and proline content means such pairs are also abundant by composition. Additionally, the attention map highlights the cationic residues R12 and R13, responsible for forming complexes with organic anions to facilitate transmembrane transport [70].

Finally, we evaluated the well-studied peptide LL-37 (**Figure S2C**). The hydrophobic N-terminus (L1 and L2) shows high relative importance, consistent with their role in driving the membrane insertion through hydrophobic interactions [58]. Concurrently, the negatively charged amino acids (D and E) reflect their experimentally established roles in driving concentration dependent aggregation of the peptides [71], [72]. These two amino acids correlate strongly across the sequence. Furthermore, the importance of the cationic K and R residues as attractors to negatively charged bacterial membranes has been discussed above, and it is also apparent here. The amino acids D and E may be used by the LL-37 to counterbalance the effect from K and R and tune selectivity against pathogenic membranes while protecting host cells [72].

Based on these results, the attention matrices successfully extract the most relevant biophysical principles learned by our generative model. Remarkably, even when averaging across multiple attention heads and selecting only one layer, distinct biological features remain preserved rather than washed out. Looking at specific heads or other layers might reveal even more granular peptide biology, which we proceed to show in “A more granular analysis at specific attention heads and layers”.

### Attention heatmap analysis on unknown peptides

**Figures 8**, **S4** and **S5** display the causal self-attention heatmaps computed for the generated peptides, whose sequences are given in Table 2. Several features stand out. A defining feature across several highly active generated peptides (P2 through P6 in **Figure 8**, and P8, P11 and P12 in **Figure S4**) is the presence of a four amino acid sequence motif (LVTG or LITG), highlighted as being important for the “AMP” property. That evidence is carried by the tag strip above each map rather than by the residue–residue map below it. This motif is absent in the wild-type (TP4) template, but it is present in other natural piscidins (piscidin-1, piscidin-2, *M. chrysops*, *M. saxatilis*) [53]. This suggests that the generative model is able to pick out entire motifs (peptide segments) with high impact on the “AMP” property and incorporate them in new sequence variants. To interpret this finding from a biophysical perspective, we need to examine two natural AMPs which contain LVTG or similar motifs (FLTG) and are among the very few that have high-resolution 3D structures determined in lipid bilayers: piscidin-1 and its homologue, piscidin-3 [8], [73]. In both cases the motif LVTG (FLTG) is helical, with L19V20 (F19L20) leading the insertion in the bilayer. The importance of the motif can be explained in light of its helicity and amphipathicity that facilitate blending into the hydrocarbon region of the bacterial membranes causing structural disruptions [8, 57]. Consistent with this direct evidence, many of our most membrane active, SI-generated constructs that carry this motif exhibit high helical contents (Tables 2 and 3). Interestingly, our CD measurements found that the P1 peptide, the most active construct of our generated set, is 100% helical, but it carries a highly charged, polyR/K segment in the C-terminus, instead of LVTG. This feature appears in the wild-type TP4 (P9 in Table 2), which ends with a polyR segment. This segment appears to be disordered according to NMR measurements [7] and corroborates our CD measurements that indicate a helical content of only 52% for P9 (Table 3). Interestingly, although P1 shows the highest helical content of all 18 peptide variants (Table 3), it also ends in RRKRSG. While polyR/K segments are, generally, expected to be disordered, it is possible that the SG at the end of P1 facilitates complete folding of the C-terminus in the bilayer environment. We learn that P1, which is somewhat surprising in its sequence architecture, embodies desirable structural features that work synergistically, for maximum action.

**Figure 8.**
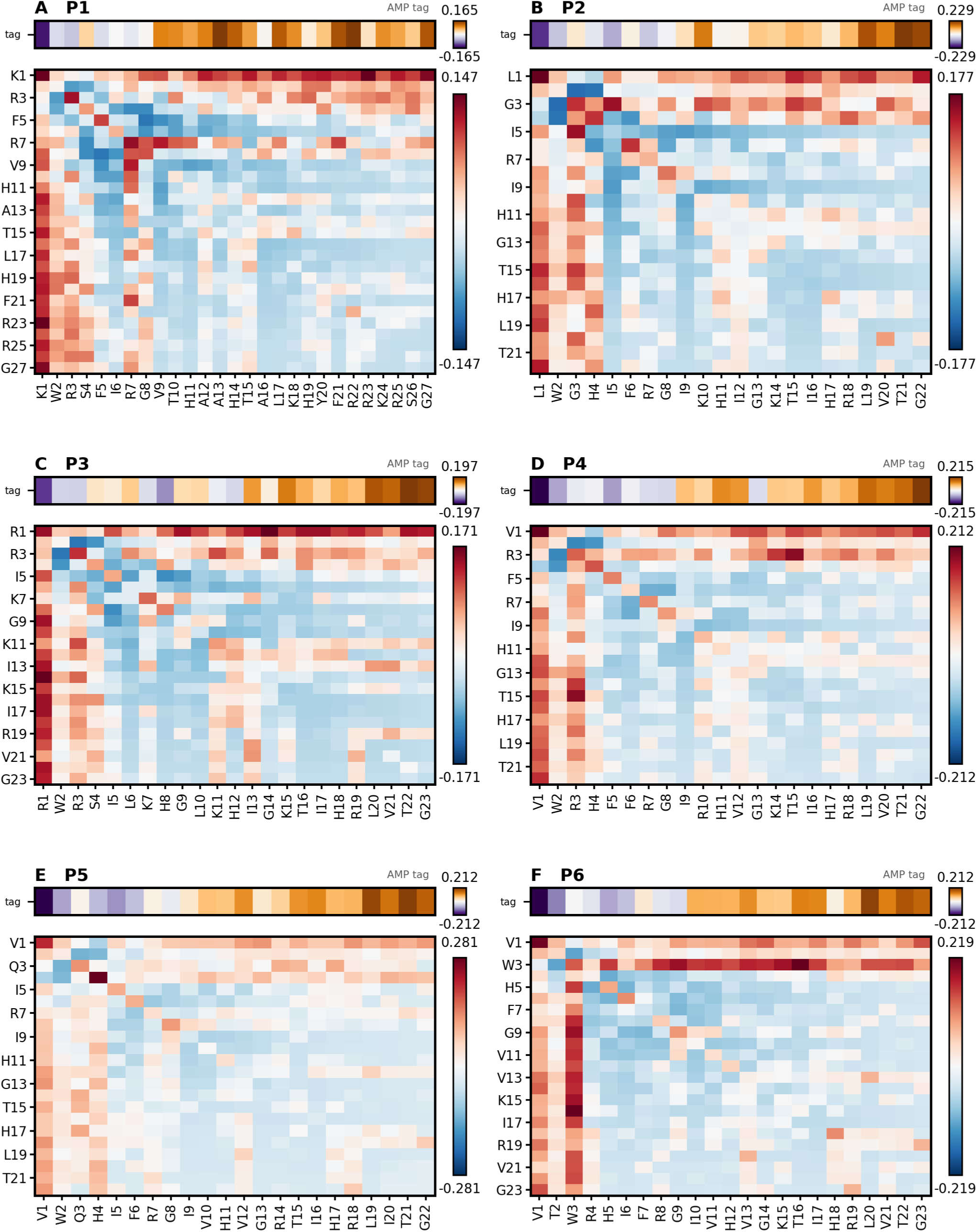
Attention heatmaps for the SI-generated peptides P1–P6 of Table 2. Combined with the experimental measurements, these heatmaps give insight into the biophysical characteristics that drive the activity of these peptides.

P2 and P3, although shorter than P1, are also very potent antimicrobials (MIC = 1 *µ*M). They feature: 1) high helical contents (77.8% and 82.5% in lipid); 2) they have an aromatic, W2 in the N-terminus, important for anchoring peptides to the membrane interface [66]; 3) they have a G13–K14 (G14–K15 in P3) and; 4) they end in the helical, amphipathic LVTG.

P4, P5 and P6 share some of the already mentioned features: a W2 (W3 for P6) at one end of the sequence, a GK (GR for P5) in the middle of the sequence, predominantly occurring in positions 13–14, and a LVTG at the other end.

Given the repetitive choice of positioning of a highly hydrophobic W2 (W3) at one end of the sequence, the LVTG (LITG) at the other end, and GK/GR in the middle of these generated constructs, we wanted to inquire further into the structural advantage for their specific placements relative to each other. A few important features that were not immediately apparent can be revealed through an examination of the helical wheel diagrams, assuming formation of helical structures. In **Figure 9** and **Figure S11** we discover that our most biologically active (lowest MIC) constructs show, besides a clear separation of hydrophobic and polar residues, a persistent alignment between the W2/W3 on the front end with another large hydrophobic residue (W2–V20, W2–L20, W2–I20, W3–V21, etc.), placed at the back end of the helix. In some cases, those pairs of aligned hydrophobes are “reinforced” by a second hydrophobic pair (V1–L19, L1–L19). It becomes apparent that these formations, protruding from the hydrophobic helical surface, may easily work together to wedge themselves between lipid headgroups, and anchor the entire peptide, irreversibly, into the membrane. This suggests that amino acids in paired positions, even if far from each other, can co-dependently influence the peptide’s action at bacterial membranes. Interestingly, P1 appears to stand apart as it displays two phenylalanine residues (F5, F21) intruding, somewhat, into the polar surface (**Figure 9A**).

**Figure 9.**
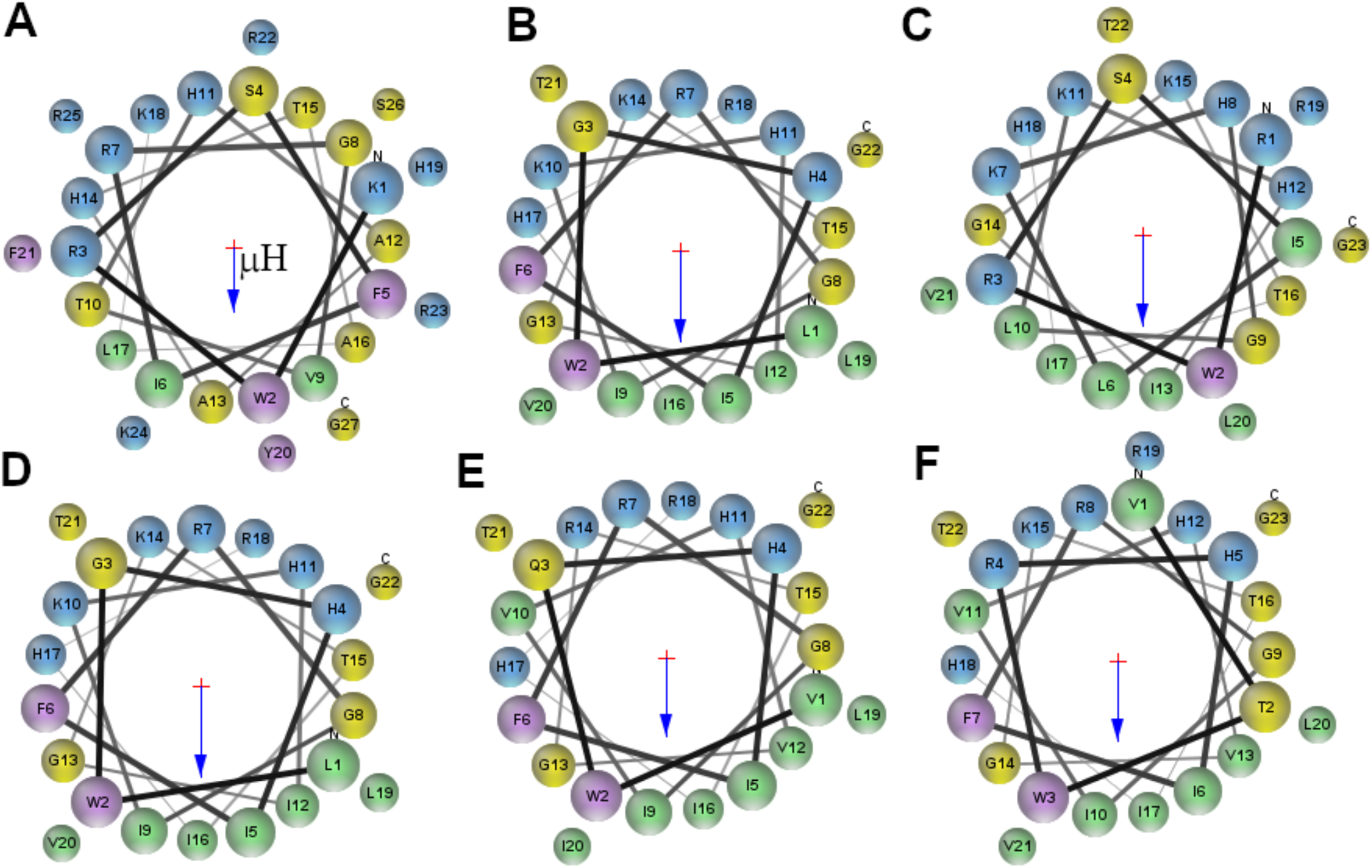
Helical wheel diagrams for the XAMPGen peptides P1–P6, shown as (A) P1, (B) P2, (C) P3, (D) P4, (E) P5 and (F) P6. The diagrams were constructed with MPEx [74], assuming the peptides form full helices in bilayers. Note that, except for P1 which shows 100% helicity in our CD measurements (Table 3), the other peptides form only partial helices with helical contents spanning a range from 64% (P6) to 83% (P3). The blue arrows indicate the hydrophobic moment vector, pointing into the direction of a (presumed) lipid bilayer surface.

Next, the GK pair is present in fourteen of the eighteen generated sequences, mostly appearing as part of a GKT triad. The same position is occupied by the GKA in the wild-type TP4, Magainin-II (**Figure 7B**) and other natural AMPs [53] [75]. The GKA motif is highly conserved in the dermaseptin family [75]. Obviously, this is a highly conserved motif, where the middle G, with its small size, is thought to form a “hinge” that imparts flexibility to the helical structure, allowing the two halves of the peptide to adjust their conformation in the membrane [8]. The following K (R in some cases) is, probably, important for maintaining the hinge tightly bound to the bacterial membrane. It is interesting that this GK pair occurs in the conserved 13–14 positions in many AMPs from various species. These positions correspond to the middle of a typical AMP sequence, whose lengths’ average is close to 26 [76]. The GKT triad appears also in the Walker A motif [77], also known as the P-loop or phosphate-binding loop [78], whose consensus G-x-x-x-x-G-K-[S/T] ends in GKT or GKS. The motif is not prominent in the attention heatmaps shown in **Figures 8**, **S4** and **S5**, because those maps report a single layer with its heads averaged and both choices suppress a feature carried by particular heads; we therefore return to it in the section below, where the head-resolved maps are examined.

Of the remaining peptides, shown in the Supporting Information, one comparison is worth making. P18 is a rationally-designed triple mutant (G7W/G8I/K14R) of P9, and it is both the weakest peptide (MIC = 64 *µ*mol/L) and the weakest binder of the entire set, with the highest *k_D_* (22.75) and the lowest *I*_max_ (2.72) (Table 3). Disrupting the two glycine linkages therefore impairs membrane binding, which supports the flexibility argument made above for the glycine residues. Also, the helical wheel diagram suggests a poor amphiphilicity associated with this peptide (**Figure S11I**).

### A more granular analysis at specific attention heads and layers

Four of the eight known peptides we examined and thirteen of our eighteen measured peptides carry a G–K/R–A/T triad, so it is worth asking where in the model that motif is actually attended to. Every attention map shown so far averages the sixteen heads of layer 22. That is a fair summary of a whole sequence, but it hides short motifs: if a single head attends strongly to three residues while the other fifteen do not, the relative importance of those residues is lost in the averaging process. The triad is a clear case. In the layer-22 maps with the heads averaged, which are the maps used in all the heatmaps displayed until now, the triad appears less important than an ordinary residue. When the maps are resolved head by head and layer by layer, the triad shows up and the signal is shown to be concentrated in a single head. Table 4 lists, for every occurrence of the GKT motif, the single layer and head that attends to it most strongly: head 11 of layer 20 wins seven of the seventeen, and head 11 accounts for eleven of them once layers 24 and 15 are included. **Figures S6**–**S8** show those maps for wild-type TP4 and the two rationally designed mutants, Magainin-2 and the set of generated peptides carrying the motif, processed with the same transform as the earlier attention heatmaps but with the head axis left intact.

**Table 4.**
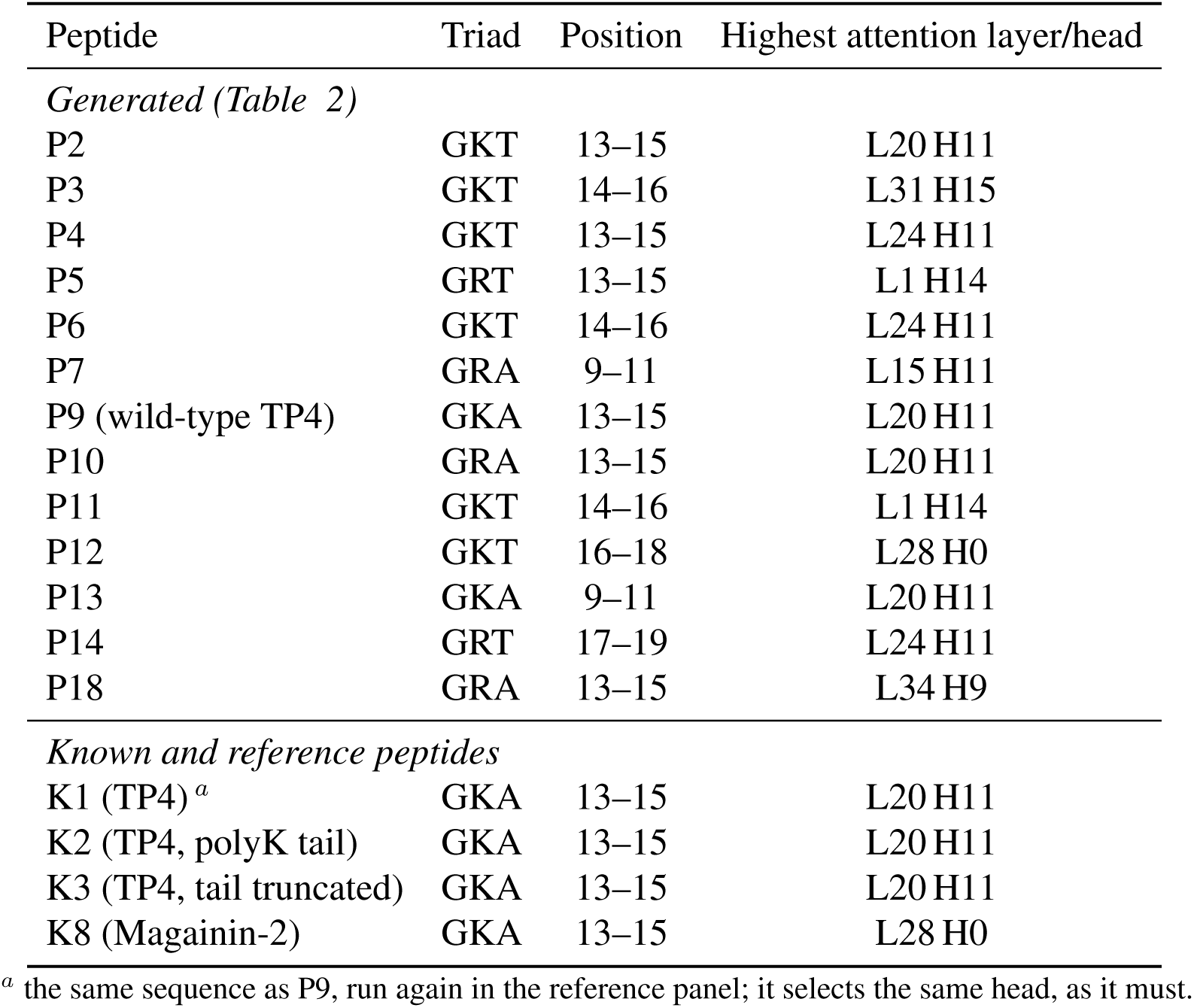
The layer and head that attends most strongly to the middle “GK”, for every occurrence of the motif in the measured and reference peptides. Each peptide is scored on all 36 16 = 576 heads and takes its own maximum. Head 11 wins eleven of the seventeen occurrences, seven of them at layer 20.

| Peptide | Triad | Position | Highest attention layer/head |
|---|---|---|---|
| *Generated (Table 2)* |  |  |  |
| P2 | GKT | 13–15 | L20 H11 |
| P3 | GKT | 14–16 | L31 H15 |
| P4 | GKT | 13–15 | L24 H11 |
| P5 | GRT | 13–15 | L1 H14 |
| P6 | GKT | 14–16 | L24 H11 |
| P7 | GRA | 9–11 | L15 H11 |
| P9 (wild-type TP4) | GKA | 13–15 | L20 H11 |
| P10 | GRA | 13–15 | L20 H11 |
| P11 | GKT | 14–16 | L1 H14 |
| P12 | GKT | 16–18 | L28 H0 |
| P13 | GKA | 9–11 | L20 H11 |
| P14 | GRT | 17–19 | L24 H11 |
| P18 | GRA | 13–15 | L34 H9 |
| *Known and reference peptides* |  |  |  |
| K1 (TP4)<sup>*a*</sup> | GKA | 13–15 | L20 H11 |
| K2 (TP4, polyK tail) | GKA | 13–15 | L20 H11 |
| K3 (TP4, tail truncated) | GKA | 13–15 | L20 H11 |
| K8 (Magainin-2) | GKA | 13–15 | L28 H0 |

<sup>*a*</sup> the same sequence as P9, run again in the reference panel; it selects the same head, as it must.

What that head is reading is the G, not the charge (K/R) following it. In the maps it is the row and column of the triad glycine that carry the coupling. The generated sequences indicate the same thing: across the peptides of Table 2 that carry the motif the model writes GKA, GKT, GRA and GRT interchangeably, varying the cation and the residue after it, but never altering the glycine. The sequences of known AMPs support this idea. Wild-type TP4 and its poly-lysine variant select the same head (Table 4), so the triad computation is independent of the polyR tail. This is consistent with our earlier conclusion that the tail acts as a separate functional module. Magainin-2, a natural peptide from an unrelated family, carries GKA at the same position as wild-type TP4. The analyzed head also treats the glycine as more than a local feature: in almost every peptide it is decoupled from every residue upstream of it and strongly coupled to every residue downstream. The model is conserving this important glycine residue, which, as mentioned above, acts as a hinge between the hydrophobic N-terminal helix and the C-terminal LVTG/polyR-carrying segment, conferring conformational flexibility.

## Discussion

Antimicrobial peptides have received a flurry of attention within the last decade, triggered by both the need to find alternative means for combating AMR, and the fast developments in computational tools and resources, especially SI tools and LLMs in the protein domain (PLMs). These tools boast immense predictive power, which stems from their ability to analyze complex datasets and uncover features and patterns that may escape human scrutiny.

LLMs in general are good generative SI tools; however, they have been received with skepticism, which originated, primarily, from the “black box” concept associated with them, wherein the decision-making process is hidden to the general user. This has led to the development of various XSI techniques to uncover the hidden processes within these LLMs [24].

In this work, we fine-tuned ProGen, a PLM, and developed a few tools that allowed us to get a glimpse inside the multiple layers of the decision process, using AMPs for our test. AMPs make up an interesting domain of biological molecules that, despite their small size, present challenges for predictive SI models. Unlike the millions of structurally stable proteins used by AlphaFold to predict shapes and active sites, most AMPs have unstable, variable structures, creating a significant predictive challenge. AMPs are usually unstructured in aqueous environments and acquire secondary structures only upon interaction with lipid membranes, for thermodynamic reasons that have been addressed at length, elsewhere [79]. The cell membranes of pathogens are both the ground for folding, mostly into *α*-helices, and the majority of AMPs’ primary target, wherein they can accumulate causing membrane lysis, or translocate into the cell to disrupt intracellular targets leading to cell death. Overall, less than 20% of recorded AMP sequences have secondary structure determined, by any methods, and very few of those structures are high-resolution structures resolved in true lipid bilayers [8], [80]. This scarcity of AMP structures is due to the fact that measurements of AMPs in bilayers are difficult to perform, require specialized techniques and are, traditionally, slow. Thus, SI techniques applied to AMPs rely heavily on peptide sequences along with their antimicrobial performance, broadly quantified by parameters like MIC, for various bacterial species [32] [76].

Nevertheless, in this work we wanted to explore the ability of ProGen, when properly fine-tuned to the AMP domain, to generalize and discover patterns and properties that are consequential for the peptides’ antibacterial potency, using only AMP sequences, with no explicit 3D structural information. We produced XAMPGen, an SI-driven workflow for design of AMPs with desired properties. To circumvent the severe computational and memory constraints of traditional full LLM fine-tuning, we leveraged Low-Rank Adaptation (LoRA) [22] to fine-tune the 1.2-billion-parameter generative protein language model ProGen [13]. By training less than 1 % of the total network parameters, the fine-tuned model successfully learned to output high-fidelity candidate sequences that closely replicate the essential physico-chemical property distributions—such as length, isoelectric point, hydrophobic moment, GRAVY index, and the exact amino acid composition of natural AMPs. To narrow the generated sequence space into effective antimicrobial candidates, we routed the outputs through a multi-stage screening pipeline powered by an encoder-only ESM model [15]. Ultimately, a final cohort of 15 candidate peptides was selected for chemical synthesis based on optimized lengths, water solubility, and localized spatial proximity within the ESM representation space to the template peptide Tilapia Piscidin-4.

We found that the synthesized sequences have strong antimicrobial activity in the assays reported here. Once we have generated and experimentally validated the peptides, we proceed forward to explain the generation process through various XSI techniques.

Through layer probing across the deep transformer architecture, we mapped the internal information flow and uncovered a distinct “U-shaped” trajectory of feature representation, isolating layer 22 as a highly performant “cornerstone layer” dense in task-specific classification, regression and helicity markers. Furthermore, by implementing a novel causal self-attention normalization technique (*A_eff_*= *A_trained_ A_untrained_*), we removed the uniform causal reference to examine residual attention patterns alongside experimental observations. This effective attention matrix faithfully captured well-documented biophysical interactions across a wide variety of literature reference sequences.

After establishing our XSI technique on well documented AMPs from the literature, we moved on to use our XSI technique along with our experimental measurements to study the synthesized/generated peptides. Interestingly, the generated peptides were imparted with several sequence features that can be found in the most active, natural AMPs known to date: 1) the regular succession of hydrophobic and polar residues at positions (*i*, *i* + 4) characteristic to helical, amphipathic peptides [81]; 2) the modular structure, emphasizing hydrophobic segments that partition strongly into membranes, separated from charged segments, like polyR motifs, that drive the efficient recruitment of the AMPs to the negatively charged, pathogenic membranes [7]; 3) the helix-breaking glycine placed in the middle of the sequence that allows the two helical halves to adjust their conformation in membranes [8] ([82]). Several of these features are familiar, drawing from our previous studies of AMPs where we combined high-resolution piscidin structures in membranes with biophysical measurements to clarify their molecular mechanisms of action [8], [73]. However, the generated peptides also contained features that were not immediately obvious to us. An example is the tryptophan residue placed in position 2 (3), that appears to align, almost perfectly, with another large hydrophobic amino acid at the other end of the helix, in position 20 (21) (**Figure 9**). This duet of large aromatic-hydrophobic residues can more strongly anchor the entire peptide into the bacterial membrane, allowing it to irreversibly insert and disrupt the membrane structure. Consistent with this interpretation is the relative position of the two large hydrophobes, 18 residues apart, spanning a distance of 27 Å, the thickness of a typical lipid membrane hydrocarbon region. This physical match would facilitate the transverse insertion of the peptide into the lipid bilayer, resulting in the formation of peptide-lined pores, or defects, as a probable mechanism of action (see [33], and the references therein).

Although we chose to work within the TP4 sequence space, for the ease of testing of the XAMPGen approach, some of the generated peptides are surprisingly novel at the sequence level, both against the AMPs of the training data and against protein sequences from the literature. Tables S2 and S3 of the Supporting Information compare the generated peptides with training AMPs and Swiss-Prot sequences using BLAST alignments [83, 84, 85]. Seven of them — P2, P4, P5, P6, P8, P11 and P12 — align to a piscidin or a close variant of it at *E* 7 10*^−^*^6^, and are therefore scaffold homologues rather than novel sequences. P1, P7, P14, P15, P16 and P17 instead reach their closest training AMP only at *E* 0.02, and their closest window in the proteome only at *E* 4.8, so these six show weaker similarity to the searched reference sets, while P3 and P13 fall between the two groups, at *E* = 1 × 10*^−^*^3^ and 6 × 10*^−^*^3^.

Ultimately, these findings provide strong evidence that XAMPGen accurately captures key physical and chemical features that can be explained and interpreted in terms of AMP structure and action at the molecular level. Moving forward, we envision incorporating structural feedback from high-throughput experiments into this XSI framework, for a faster and more guided exploration of the vast peptide sequence space, in search for exceptional candidates.

## Conclusion

In this work, we introduced XAMPGen, an interpretable SI pipeline designed for the discovery of novel AMPs. Our predictive framework combines elements of PLMs (ProGen), Low-Rank Adaptation (LoRA), a deep binary classifier and a MIC regression model to evaluate and filter peptide candidates. We dove into the decision-making process to show that the model learns and internalizes features of AMP sequences that can be translated into meaningful structural attributes that are consequential for the sought property (e.g., low MIC values). Insights from the generative process and attention matrix analyses were evaluated with direct, experimental measurements of a set of XAMPGen-designed peptides in lipid membrane platforms. The experimental data revealed that XAMPGen was able to interlace structural motifs and attributes found in highly potent, natural AMPs, with other, less known features, to generate exceptional candidates, like P1. These candidates merit further investigations into their therapeutic profiles and suitability for drug development. Finally, XAMPGen addresses the persistent “black box” challenge of deep learning models in computational biology by employing XSI techniques to reveal the underlying mechanisms behind sequence generation. By combining parameter-efficient sequence generation with high-throughput biophysical measurement and validation, this framework provides a relatively fast and trustworthy toolkit to accelerate the discovery of potent AMPs and also to explore a wider variety of membrane-active peptides for medical applications.

## Code availability

The XAMPGen source code is available at https://github.com/mondalkinjal/XAMPGen.

## Acknowledgments

This work was supported by the National Institute of Standards and Technology (NIST).

## Disclaimer

Certain commercial equipment, instruments, software, or materials are identified in this paper in order to specify the experimental procedure adequately. Such identification is not intended to imply recommendation or endorsement of any product or service by the National Institute of Standards and Technology, nor is it intended to imply that the materials or equipment identified are necessarily the best available for the purpose.

