## Supporting Information for "XAMPGen: An Interpretable Super Intelligence Pipeline For Novel Antimicrobial Peptide Discovery"


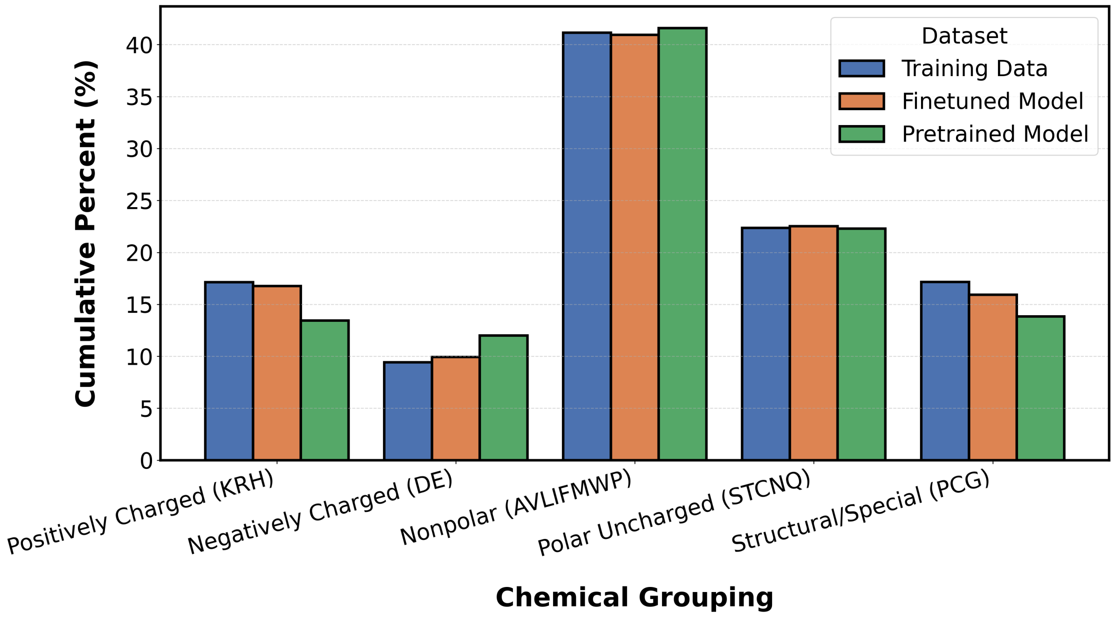


**Figure S1.** Amino acid composition, by chemical class, of the AMP training data (blue) and of 20,000 sequences each generated by ProGen after LoRA fine-tuning (orange) and by the pretrained ProGen without fine-tuning (green). Each bar is the percentage of all residues in a set that belong to the class. The fine-tuned model, unlike the pretrained model, reproduces the charge and structural/special composition of the training AMPs.

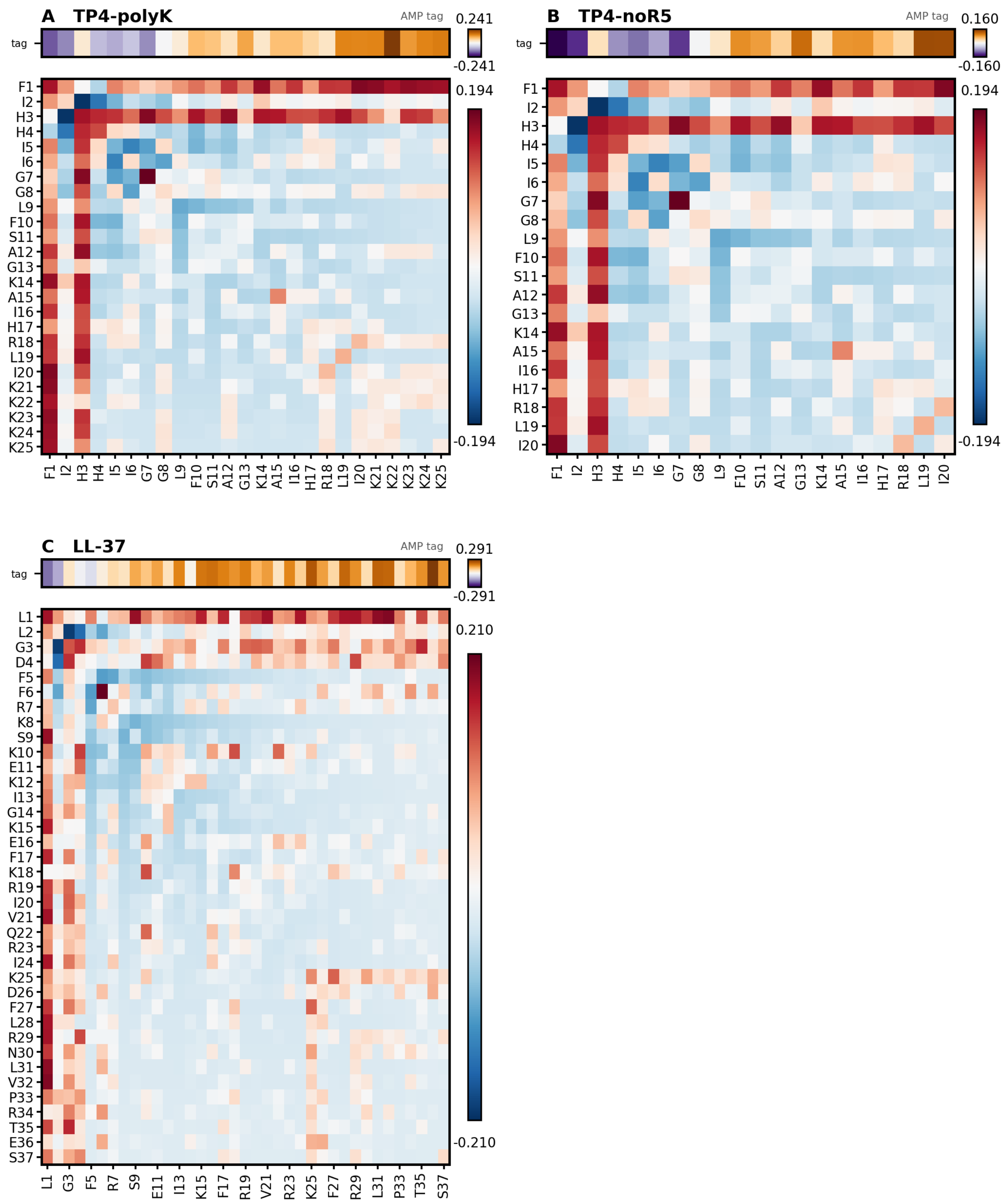


**Figure S2.** Attention heatmaps at layer 22 for TP4 variants and additional AMPs. Effective attention between token pairs — between residues, and between each residue and the antimicrobial control tag (AMP) — for (A) TP4 with the C-terminal polyR tail substituted by polyK, FIHHIIGGLFSAGKAIHRLIKKKKK; (B) TP4 truncated after I20, FIHHIIGGLFSAGKAIHRLI; (C) LL-37, LLGDFFRKSKEKIGKEFKRIVQRIKDFLRN-LVPRTES.

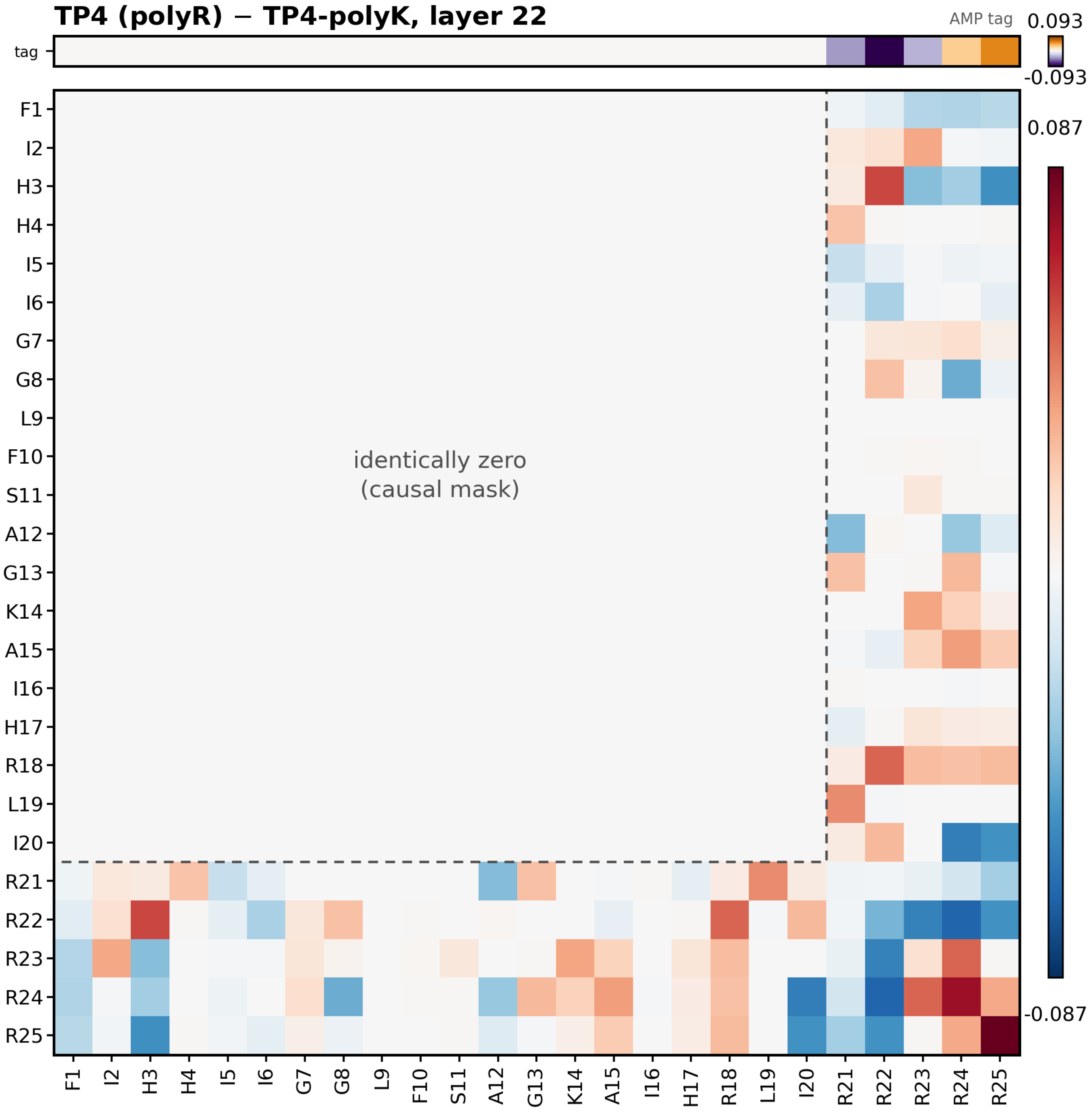


**Figure S3.** The polyR → polyK tail substitution at layer 22, read as a difference map: head-averaged effective attention of wild-type TP4 minus that of the polyK variant. Red marks a coupling stronger with polyR, blue one stronger with polyK.

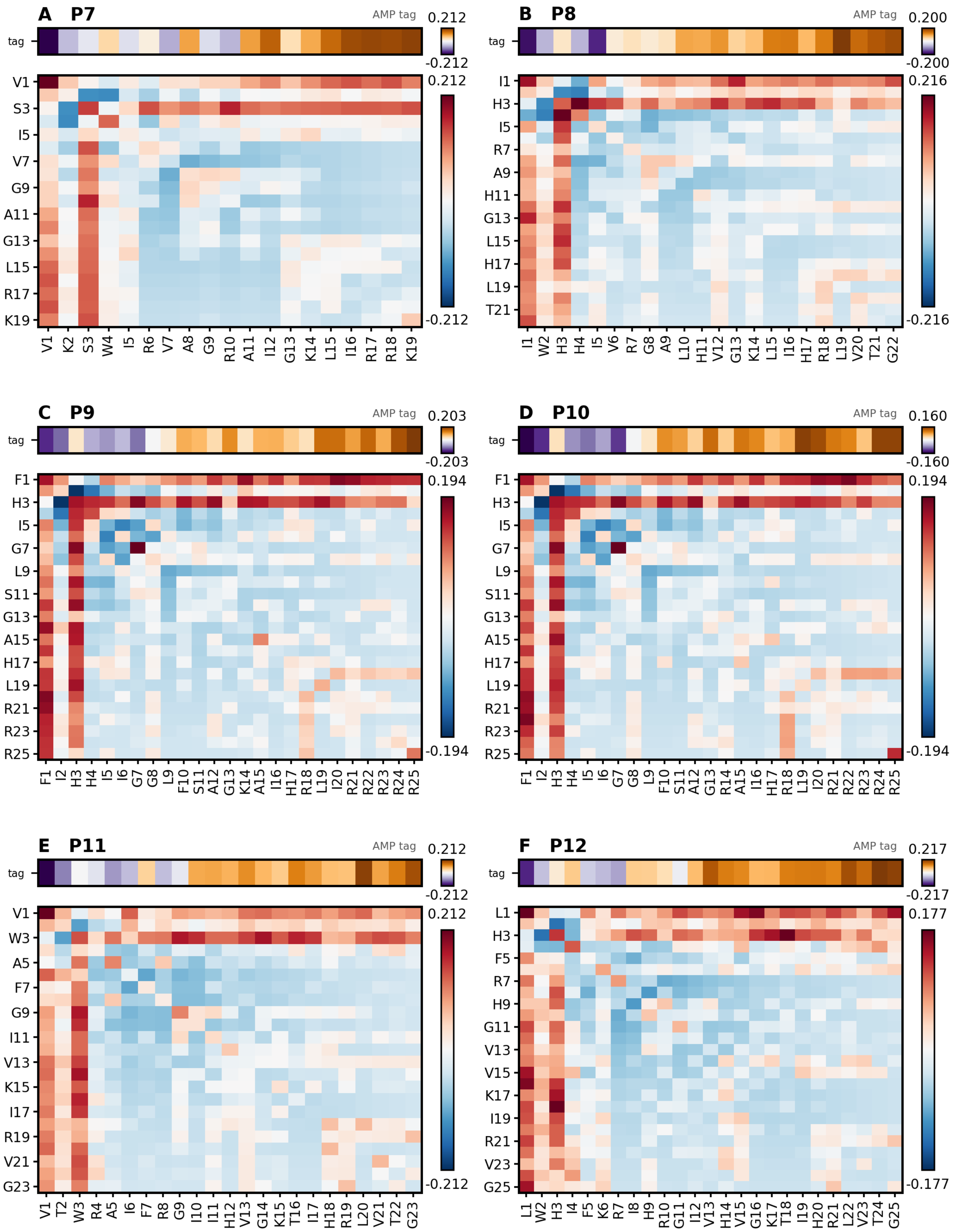


**Figure S4.** Attention heatmaps at layer 22 for unknown AMPs P7–P12 of Table 2, one of them being the wild-type TP4 peptide P9.

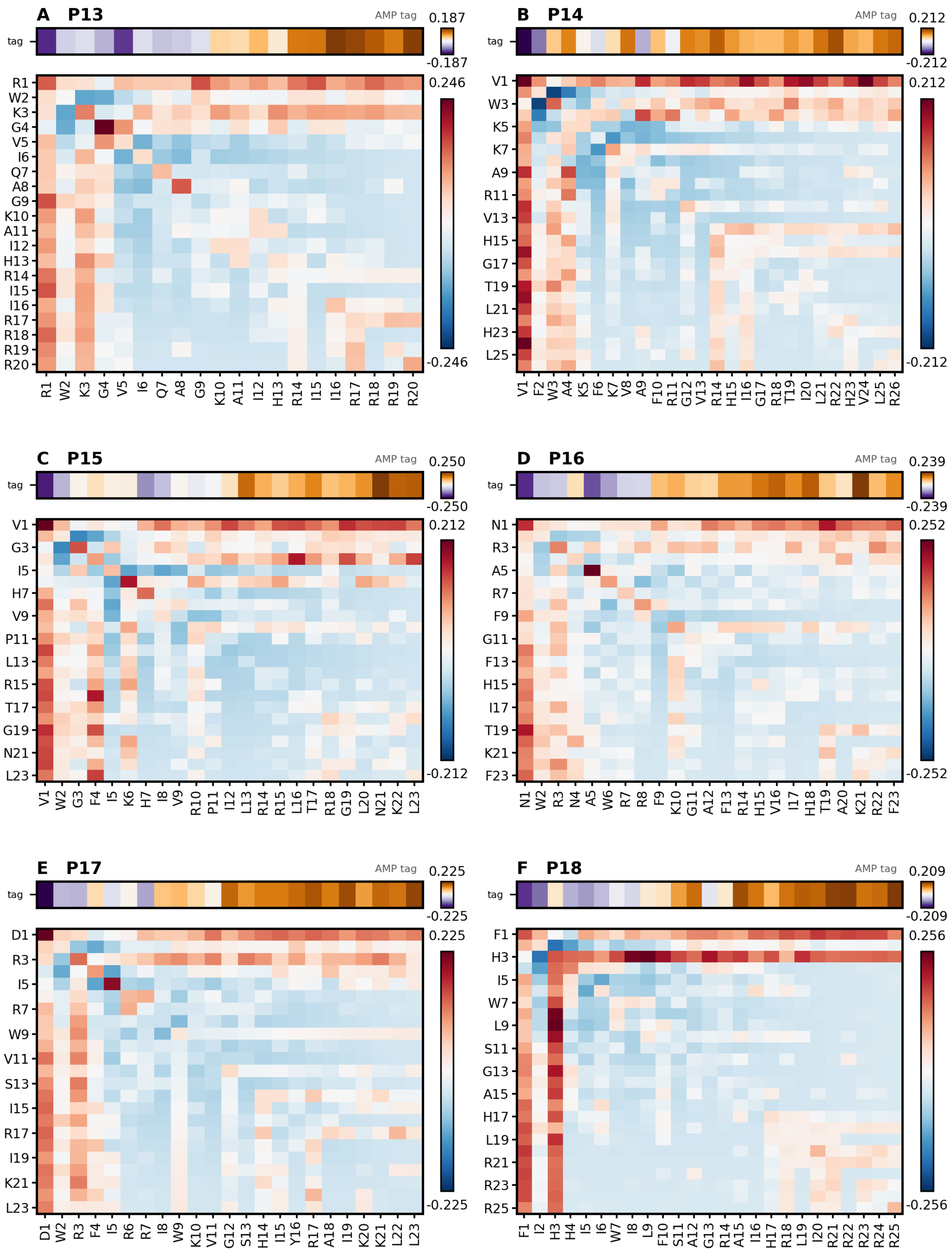


**Figure S5.** Attention heatmaps at layer 22 for unknown AMPs P13–P18 of Table 2. Drawn as in **Figure S4**.

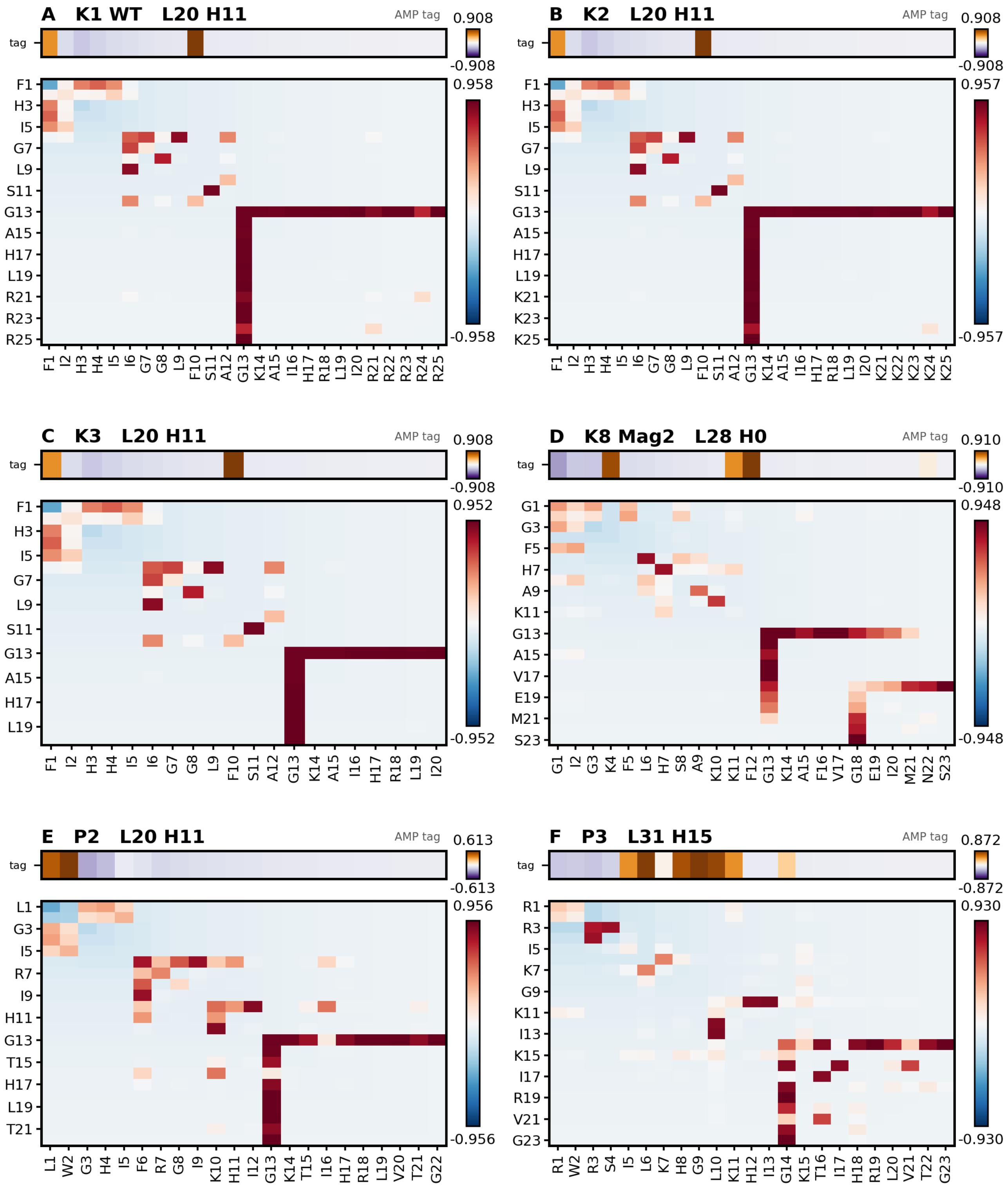


**Figure S6.** Attention paid to the G–K/R–A/T triad by the single head, out of the 36 $\times$ 16 $=$ 576 in the model, that responds to it most strongly, for the reference peptides and the first two generated ones. Each panel heading names the peptide followed by its layer and head, so that “P2, L20 H11” is peptide P2, layer 20, head 11. The layer and head shown for each peptide are those listed in Table 4 of the main text.

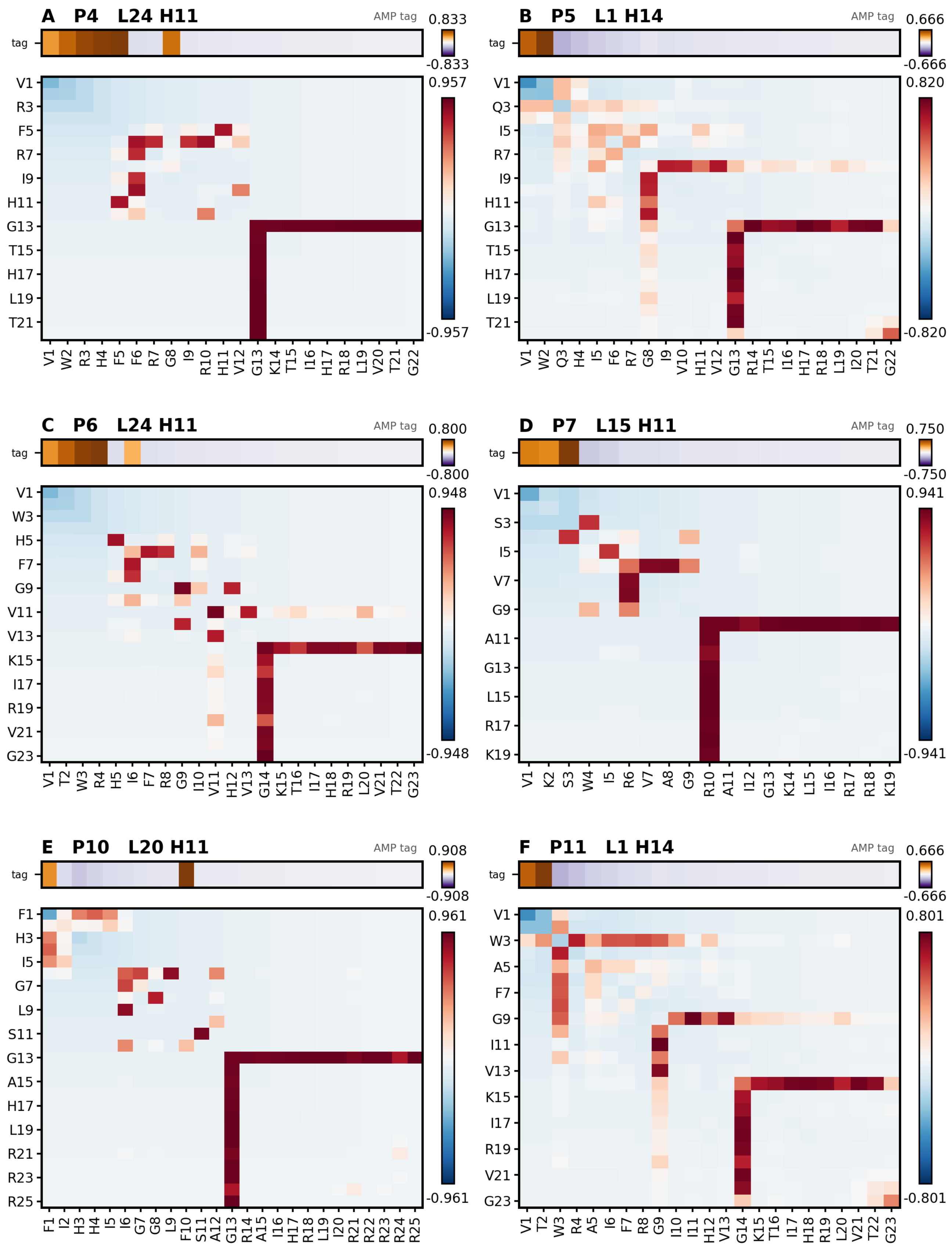


**Figure S7.** As **Figure S6**, for the generated peptides P4–P11; the heads shown are those listed in Table 4 of the main text.

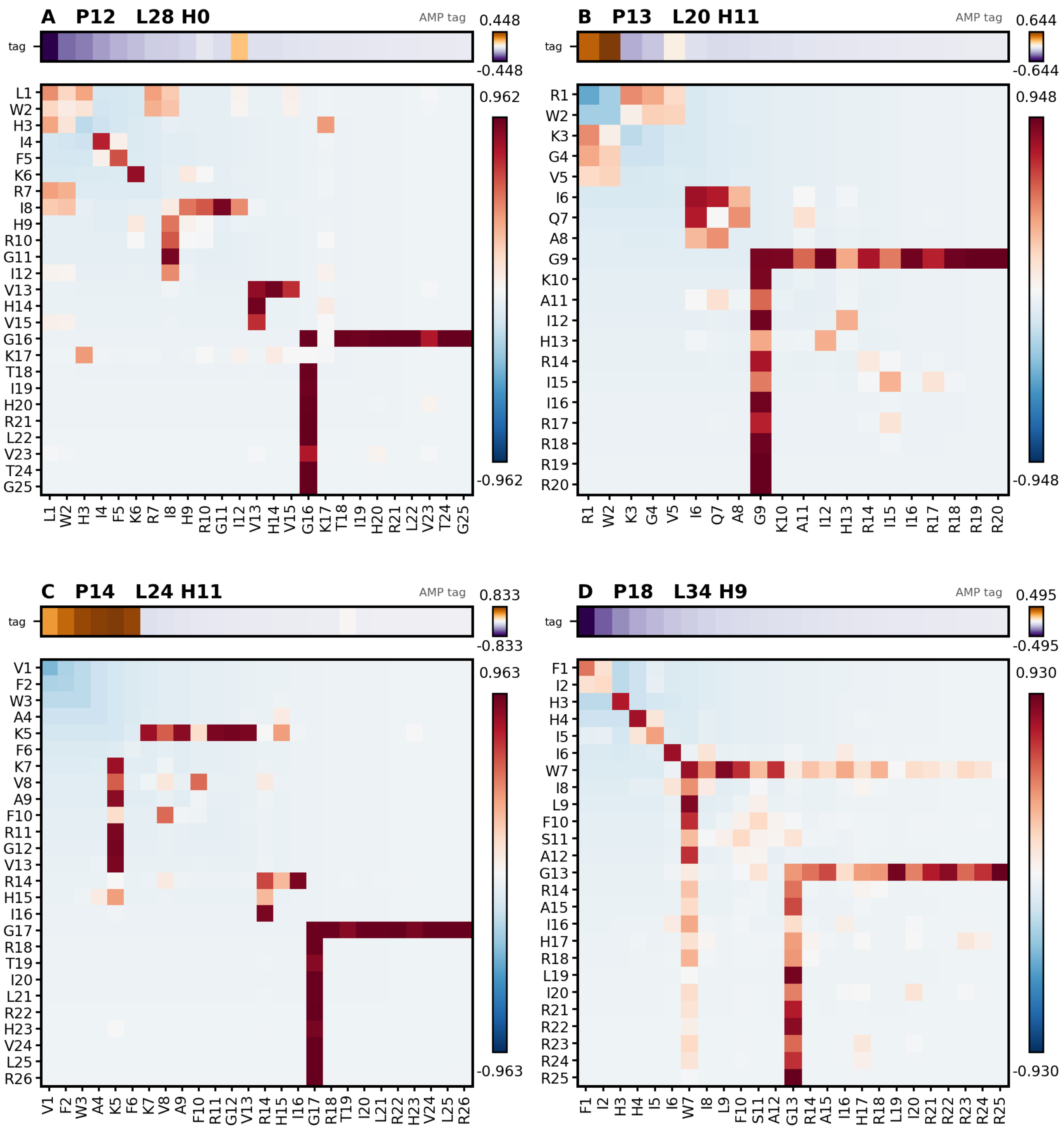


**Figure S8.** As **Figure S6**, for the remaining generated peptides; the heads shown are those listed in Table 4 of the main text.

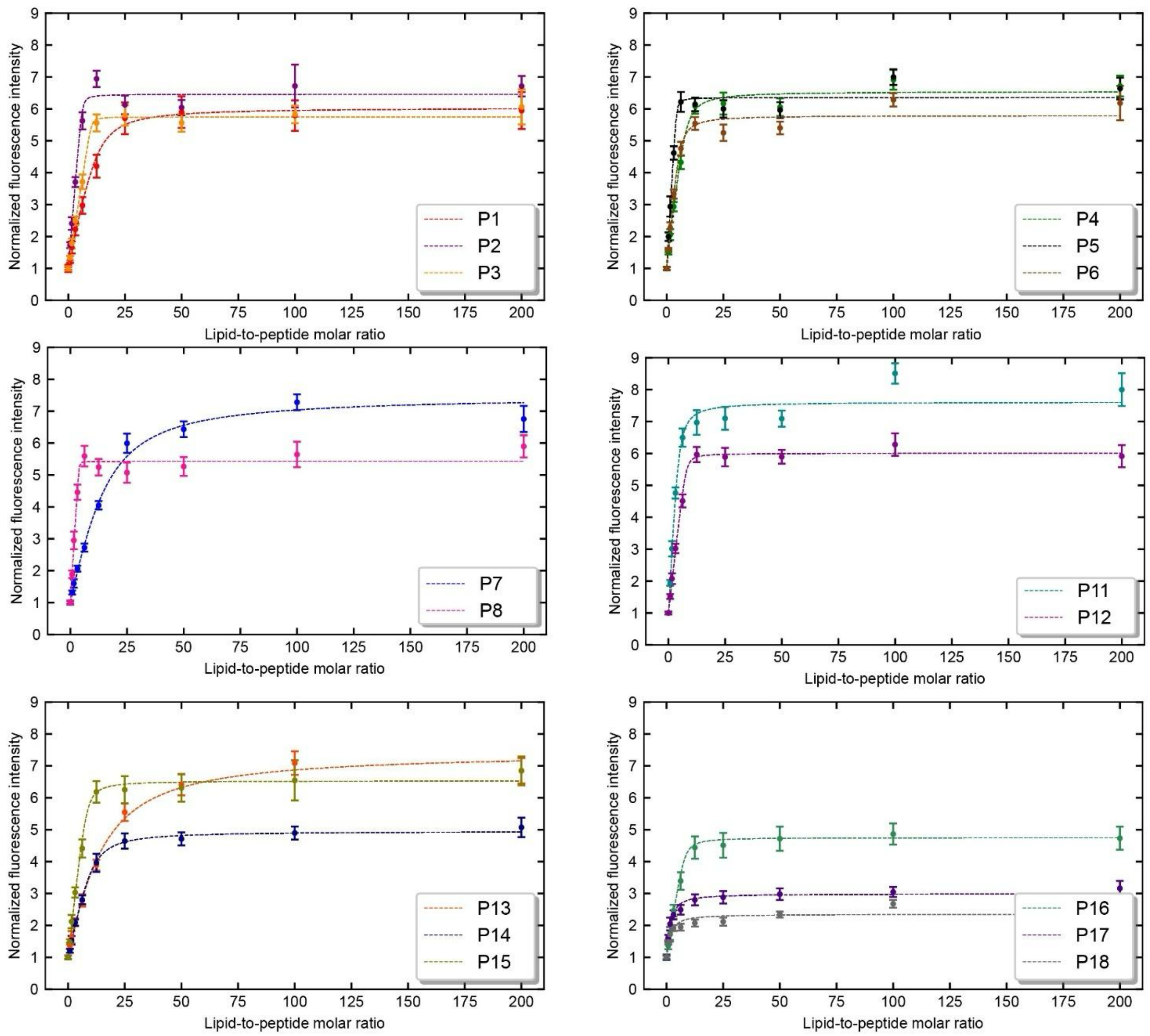


**Figure S9.** Fluorescence intensities for the peptides measured at 334 nm at various lipid-to-peptide (L/P) ratios. The intensities were normalized to the fluorescence intensity at zero lipid concentration and corrected for light scattering as described by Ladokhin et al [1]. All the samples included 10 μmol/L peptide mixed with different concentrations of small unilamellar vesicles composed of POPC/POPG (3:1 molar ratio) and containing 5 mol% DMG-PEG$_{2000}$ to achieve L/P ratios ranging from 0 to 200. Measurements were performed in duplicate, and the error bars represent the standard deviation. The resulting data were fitted to a modified Langmuir binding model using a Markov Chain Monte Carlo sampling method as described in our previous publication [2] to get fitted parameters including the dissociation constant ($K_{\mathrm{D}}$), number of lipids bound per peptide ($n$), and the maximum fluorescence intensity ($I_{\max}$).

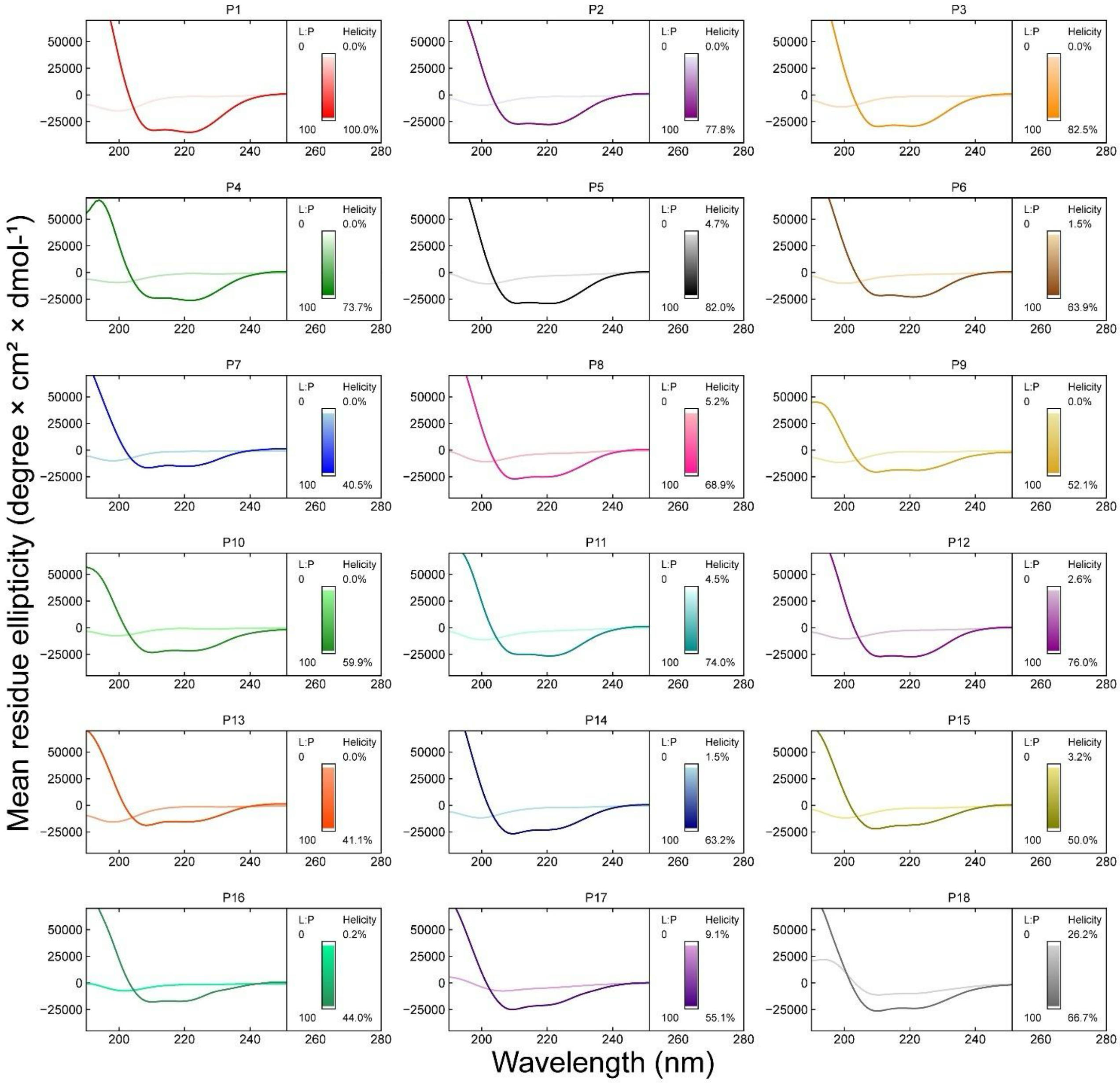


**Figure S10.** Circular dichroism (CD) spectra for the 17 generated peptides and TP4 (wildtype). Spectra show the mean residue ellipticity (MRE) of all the peptides in buffer (5 mmol/L sodium phosphate, 5 mmol/L NaCl buffer, pH 7) and in presence of small unilamellar vesicles composed of POPC/POPG (3:1 molar ratio) containing 5 mol% DMG-PEG$_{2000}$ at a lipid-to-peptide molar ratio of 100:1. All samples were prepared at 10 µmol/L peptide and 1000 µmol/L lipid and measured at 25 °C. All CD spectra were smoothened using a Gaussian fit.

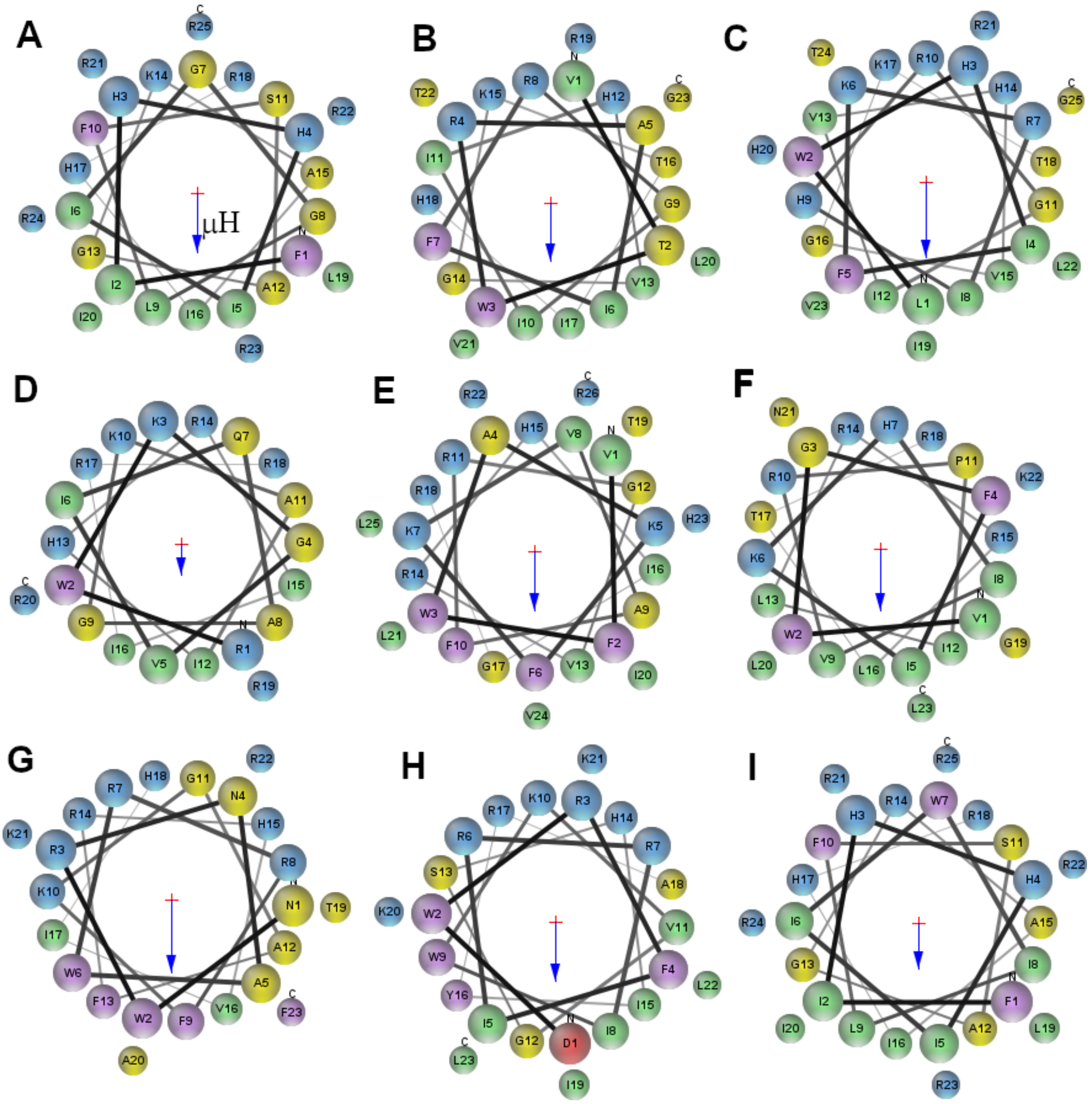


**Figure S11.** Helical wheel diagrams for XAMPGen peptides. (A) P9; (B) P11; (C–I) P12–P18. The diagrams were constructed with MPEx [3], assuming the peptides form full helices in bilayers. The blue arrows indicate the hydrophobic moment vector, pointing into the direction of a (presumed) lipid bilayer surface.

**Table S1.** Peptides sorted by *E. coli* MIC potency and numbered sequentially P1 through P18. Parameters calculated using the White and Wimley scale from the DBAASP database.

| Name | Peptide sequence | Norm $\mu$H (octanol WW) | Norm H (octanol WW) | Net Charge | Isoelectric Point | Disordered Propensity | Linear Moment | Aggregation Propensity |
|---|---|---|---|---|---|---|---|---|
| **P1** | `KWRSFIRGVTHAAHTALKHYFRRKRSG` | 0.51 | 0.76 | 8 | 12.31 | -0.36 | 0.27 | 0 |
| **P2** | `LWGHIFRGIKHIGKTIHRLVTG` | 0.88 | 0.46 | 4 | 12.15 | 0.09 | 0.24 | 0 |
| **P3** | `RWRSILKHGLKHIGKTIHRLVTG` | 0.84 | 0.68 | 6 | 12.43 | -0.15 | 0.25 | 0 |
| **P4** | `VWRHFFRGIRHVGKTIHRLVTG` | 0.87 | 0.48 | 5 | 12.58 | -0.04 | 0.25 | 0 |
| **P5** | `VWQHIFRGIVHVGRTIHRLITG` | 0.76 | 0.28 | 3 | 12.4 | 0.21 | 0.25 | 0 |
| **P6** | `VTWRHIFRGIVHVGKTIHRLVTG` | 0.74 | 0.4 | 4 | 12.41 | 0.14 | 0.21 | 0 |
| **P7** | `VKSWIRVAGRAIGKLIRRK` | 0.55 | 0.62 | 7 | 12.6 | -0.17 | 0.25 | 0 |
| **P8** | `IWHHIVRGALHVGKLIHRLVTG` | 0.71 | 0.43 | 3 | 12.12 | 0.26 | 0.23 | 0 |
| **P9** | `FIHHIIGGLFSAGKAIHRLIRRRRR` | 0.64 | 0.56 | 7 | 12.8 | -0.08 | 0.23 | 27.19 |
| **P10** | `FIHHIIGGLFSAGRAIHRLIRRRRR` | 0.61 | 0.52 | 7 | 12.88 | -0.09 | 0.23 | 27.12 |
| **P11** | `VTWRAIFRGIIHVGKTIHRLVTG` | 0.67 | 0.29 | 4 | 12.41 | 0.22 | 0.24 | 0 |
| **P12** | `LWHIFKRIHRGIVHVGKTIHRLVTG` | 0.81 | 0.49 | 5 | 12.42 | 0.08 | 0.27 | 0 |
| **P13** | `RWKGVIQAGKAIHRIIRRRR` | 0.43 | 0.79 | 8 | 12.81 | -0.43 | 0.21 | 0 |
| **P14** | `VFWAKFKVAFRGVRHIGRTILRHVLR` | 0.61 | 0.35 | 7 | 12.71 | 0.05 | 0.19 | 0 |
| **P15** | `VWGFIKHIVRPILRRLTRGLNKL` | 0.72 | 0.24 | 6 | 12.59 | 0.03 | 0.22 | 0 |
| **P16** | `NWRNAWRRFKGAFRHVIHTAKRF` | 0.85 | 0.57 | 7 | 12.71 | -0.39 | 0.23 | 0 |
| **P17** | `DWRFIRRIWKVGSHIYRAIKKLL` | 0.69 | 0.42 | 6 | 11.76 | -0.1 | 0.19 | 0 |
| **P18** | `FIHHIIWILFSAGRAIHRLIRRRRR` | 0.5 | 0.3 | 7 | 12.88 | -0.03 | 0.3 | 603.65 |

### Sequence novelty of the generated peptides

**Table S2.** Closest sequence to each measured peptide among the 22,826 AMPs of the fine-tuning set, by BLASTP (`blastp-short`); the self-match is excluded for P9. Residues in **bold** are aligned identically, and "% id." is the number of such residues divided by the length of the measured peptide. Entries marked † are precursors, for which only the aligned segment is shown, with the precursor's full length in parentheses.

| Peptide | Origin | Sequence | Closest sequence in the fine-tuning set | % id. | $E$ |
|---|---|---|---|---|---|
| P9 | wild type | **FIHHIIGGLFSAGKAIHRLIRRR**RR | **FIHHIIGGLFSAGKAIHRLIRRR** | 92.0 | $2 \times 10^{-9}$ |
| P10 | human | **FIHHIIGGLFSAG**R**AIHRLIRRR**RR | **FIHHIIGGLFSAG**K**AIHRLIRRR** | 88.0 | $7 \times 10^{-9}$ |
| P18 | human | **FIHHII**WI**LFSAG**R**AIHRLIRRR**RR | **FIHHII**GG**LFSAG**K**AIHRLIRRR** | 80.0 | $2 \times 10^{-6}$ |
| P6 | generated | VT**W**R**HIFRGIVHVGKTIHRLVTG** | F**W**H**HIFRGIVHVGKTIHRLVTG** | 87.0 | $2 \times 10^{-8}$ |
| P2 | generated | LWG**HIFRGIKH**I**GKTIHRLVTG** | FFH**HIFRGIKH**V**GKTIHRLVTG** | 81.8 | $7 \times 10^{-7}$ |
| P4 | generated | V**W**R**H**F**FRGI**R**HVGKTIHRLVTG** | F**W**H**H**I**FRGI**V**HVGKTIHRLVTG** | 81.8 | $1 \times 10^{-6}$ |
| P5 | generated | V**W**Q**HIFRGIVHVG**R**TIHRL**I**TG** | F**W**H**HIFRGIVHVG**K**TIHRL**V**TG** | 81.8 | $1 \times 10^{-7}$ |
| P11 | generated | VT**W**RA**IFRGI**I**HVGKTIHRLVTG** | F**W**HH**IFRGI**V**HVGKTIHRLVTG** | 78.3 | $4 \times 10^{-7}$ |
| P8 | generated | IW**HHI**V**RGA**L**HVGK**L**IHRLVTG** | FF**HHI**F**RGA**V**HVGK**T**IHRLVTG** | 77.3 | $7 \times 10^{-6}$ |
| P12 | generated | LWH**IF**KR**I**HR**GIVHVGKTIHRLVTG** | **IF**HH**I**FK**GIVHVGKTIHRLVTG**† (67) | 72.0 | $4 \times 10^{-6}$ |
| P13 | generated | RW**KG**V**I**Q**AGKAIH**RI**I**R**RRR** | FFGWLI**KG**A**I**H**AGKAIH**PL**I**H**RRR**H | 65.0 | $6 \times 10^{-3}$ |
| P3 | generated | RWRSIL**K**H**G**L**KH**I**GKTIHRLVTG** | FFHHI**K**R**G**I**KH**V**GKTIHRLVTG** | 60.9 | $1 \times 10^{-3}$ |
| P17 | generated | DWRF**I**R**R**I**W**K**VG**SHI**YRAIKK**LL | **I**K**R**F**W**P**V**VIRTVVA**G**YNL**YRAIKK**† (148) | 47.8 | $2 \times 10^{-2}$ |
| P7 | generated | VKS**W**I**R**VA**GRAIG**KL**IR**RK | **W**L**R**KT**GRAIG**NA**IR**† (162) | 47.4 | $4 \times 10^{-2}$ |
| P16 | generated | NWRN**AWR**RF**K**G**A**F**R**H**V**IH**TAK**RF | GWFKK**AWR**KV**K**H**A**G**R**R**V**LD**TAK**GVGRHYLNNWLNRYRG | 43.5 | 3.5 |
| P14 | generated | VFWA**KF**KVA**FRG**VR**H**I**G**R**TI**L**R**H**V**LR | **KF**HHI**FRG**IK**H**V**G**K**TI**H**R**L**V**TG | 42.3 | 0.27 |
| P1 | generated | **KW**R**SF**I**R**GVTHA**AHTALK**HYFRRKRSG | **KW**K**SF**L**R**TLKSP**A**KTVF**HTALK**AISS | 40.7 | 5.6 |
| P15 | generated | VW**G**FI**K**H**I**V**RP**I**LRR**LTRGLNKL | KILR**G**VC**K**K**I**M**RP**F**LRR**ISKDILTGKK | 34.8 | 5.2 |

**Table S3.** Closest window in UniProtKB/Swiss-Prot [4] for each measured peptide, by the same search. Residues in **bold** are aligned identically, and "% id." is again divided by the length of the measured peptide. Each window is given with its residue range and the protein's full length in parentheses.

| Peptide | Sequence | Closest window in Swiss-Prot | Protein | % id. | $E$ |
|---|---|---|---|---|---|
| P9 | **FIHHIIGGLF**SA**GK**A**IH**R**LIR**RRR**R** | **FIHHIIGGLF**HV**GK**S**IH**D**LIR**GKN**R** | Pteroicidin-α (23–47 of 66) | 72.0 | $2 \times 10^{-3}$ |
| P10 | **FIHHIIGGLF**SA**G**RA**IH**R**LIR**RRR**R** | **FIHHIIGGLF**HV**G**KS**IH**D**LIR**GKN**R** | Pteroicidin-α (23–47 of 66) | 68.0 | $6 \times 10^{-3}$ |
| P18 | **FIHHII**WI**LF**SA**G**RA**IH**R**LIR**RRR**R** | **FIHHII**GG**LF**HV**G**KS**IH**D**LIR**GKN**R** | Pteroicidin-α (23–47 of 66) | 60.0 | 0.94 |
| P6 | VTWR**HIFRGIVHVGKTIHRLVTG** | FH**HIFRGIVHVGKTIHRLVTG** | Moronecidin (24–44 of 79) | 82.6 | $4 \times 10^{-5}$ |
| P2 | LWG**HIFRGI**K**H**I**GKTIHRLVTG** | **HIFRGI**V**H**V**GKTIHRLVTG** | Moronecidin (26–44 of 79) | 77.3 | $2 \times 10^{-3}$ |
| P4 | VWR**H**F**FRGI**R**HVGKTIHRLVTG** | FH**H**I**FRGI**V**HVGKTIHRLVTG** | Moronecidin (24–44 of 79) | 77.3 | $2 \times 10^{-3}$ |
| P5 | VWQ**HIFRGIVHVG**R**TIHRL**I**TG** | FH**HIFRGIVHVG**K**TIHRL**V**TG** | Moronecidin (24–44 of 79) | 77.3 | $1 \times 10^{-4}$ |
| P11 | VTWRA**IFRGI**I**HVGKTIHRLVTG** | FHH**IFRGI**V**HVGKTIHRLVTG** | Moronecidin (24–44 of 79) | 73.9 | $1 \times 10^{-3}$ |
| P8 | IW**HHI**V**RG**AL**HVGK**L**IHRLVTG** | FF**HHI**F**RG**IV**HVGK**T**IHRLVTG** | Moronecidin (23–44 of 79) | 72.7 | $1 \times 10^{-2}$ |
| P12 | LWHI**F**KR**I**H**RGIVHVGKTIHRLVTG** | F**F**HH**I**F**RGIVHVGKTIHRLVTG** | Moronecidin (23–44 of 79) | 72.0 | $2 \times 10^{-3}$ |
| P13 | RW**KG**V**I**Q**AGKAIH**RI**I**R**RRR** | **KG**A**I**H**AGKAIH**GL**I**H**RRR** | Chrysophsin-1 (7–24 of 25) | 65.0 | 3.5 |
| P3 | R**WRSILKHGLK**H**I**G**KTI**HRLVTG | **WRSIL**CKEAR**KH**DS**GLK**LFKLGIKLCTRNEQ**I**M**KTI** | Mediator of RNA pol. II subunit (838–873 of 1148) | 60.9 | 11 |
| P17 | DW**RF**I**RR**I**WKV**GS**HI**YRAIKKLL | **RF**L**RR**L**WK**A**V**EG**HI** | Leucine–tRNA ligase (671–684 of 864) | 39.1 | 4.8 |
| P7 | VKSW**IR**V**AGRA**I**G**K**LIRR**K | **IR**T**AGRA**A**G**I**L**ALDDAT**IRR** | Pyruvate aldolase (205–224 of 257) | 57.9 | 31 |
| P16 | NWRNA**WRRF**K**GAFRHVI**HT**AK**RF | **WRR**DLQPYPSHGGR**F**T**G**LP**A**Y**FRHVI**SA**AK** | HMG-CoA synthase (185–214 of 343) | 56.5 | 11 |
| P14 | VFWAKFKVA**FR**G**VR**H**IGRTI**L**R**H**VLR** | **FRVR**LTHSLEVAQ**IGRTI**A**RVLR** | dGTP triphosphohydrolase (65–87 of 386) | 50.0 | 15 |
| P1 | KW**RSF**IR**G**V**T**H**AA**HT**AL**KHY**F**R**RKRS**G | **RSF**F**G**S**T**NSCPI**AA**NP**ALF**K**RKRS** | LRR receptor-like kinase (426–449 of 853) | 51.9 | 12 |
| P15 | VWG**FI**K**HIV**RPI**LRRLTR**G**L**N**KL** | **FI**T**HIV**KSV**L**E**RR**RDD**L**K**T**IE**R**D**L**S**KL** | Asparagine–tRNA ligase (248–274 of 442) | 60.9 | 11 |
